# Mitochondrial variation in *Anopheles gambiae* and *An. coluzzii*: phylogeographic legacy of species isolation and mito-nuclear associations with metabolic resistance to pathogens and insecticides

**DOI:** 10.1101/2023.07.18.549472

**Authors:** Jorge E. Amaya Romero, Clothilde Chenal, Yacine Ben Chehida, Alistair Miles, Chris S. Clarkson, Vincent Pedergnana, Bregje Wertheim, Michael C. Fontaine

## Abstract

Mitochondrial DNA (mtDNA) has been a popular marker in phylogeography, phylogeny, and molecular ecology, but its complex evolution is increasingly recognized. Here, we investigated mtDNA variation in *An. gambiae* and *An. coluzzii*, in perspective with other species in the *Anopheles gambiae* complex (AGC), by assembling the mitogenomes of 1219 mosquitoes across Africa. The mtDNA phylogeny of the AGC was consistent with a previously reported highly reticulated evolutionary history, revealing important discordances with the species tree. The three most widespread species (*An. gambiae, An. coluzzii, An. arabiensis*), known for extensive historical introgression, could not be discriminated based on mitogenomes. Furthermore, a monophyletic clustering of the three salt-water tolerant species (*An. merus, An. melas, An. bwambae*) in the AGC also suggested that introgression and possibly selection shaped mtDNA evolution. MtDNA variation in *An. gambiae* and *An. coluzzii* across Africa revealed significant partitioning among populations and species. A peculiar mtDNA lineage found predominantly in *An. coluzzii* and in the hybrid taxon of the African “*far-west”* exhibited divergence comparable to the inter-species divergence in the AGC, with a geographic distribution matching closely *An. coluzzii*’s geographic range. This phylogeographic relict of the *An. coluzzii* and *An. gambiae* split was associated with population and species structuration, but not with *Wolbachia* occurrence. The lineage was significantly associated with SNPs in the nuclear genome, particularly in genes associated with pathogen and insecticide resistance. These findings underline the mito-nuclear coevolution history and the role played by mitochondria in shaping metabolic responses to pathogens and insecticide in *Anopheles*.

## Introduction

Historically, mitochondrial DNA (mtDNA) has been among the most popular genetic markers in molecular ecology, evolution, and systematics. Among its applications are assessing population and species genetic diversity, genetic structure, phylogeographic and phylogenetic patterns, species identity and meta-barcoding (Galtier, et al. 2009; Dong, et al. 2021; Dowling and Wolff 2023). Contributing factors to such popularity include an easy access to the mtDNA genetic variation compared to nuclear markers, even in degraded tissue samples, due to the large number of per-cell copies. Likewise, its haploid nature and clonal inheritance through the female germ line provide an account of evolution independent, and complimentary, to the nuclear DNA’s (nuDNA). Because the mtDNA does not recombine, the entire molecule behaves as a single segregating locus, with a single genealogical tree representative of the maternal genealogy. Furthermore, its reduced effective population size together with an elevated mutation rate compared to the nuclear genome makes the mtDNA a fast-evolving, and potentially highly informative genetic marker (Galtier, et al. 2009; Allio, et al. 2017; Dong, et al. 2021; Dowling and Wolff 2023). At the same time, however, the fact that mtDNA is a single non-recombining locus limits its power to describe the evolutionary history of populations and species.

Another argument in favor of mtDNA’s popularity as a genetic marker is its near-neutrality and constant mutation rate, but increasing number of studies now contend that selection and other factors can significantly impact mtDNA variation and its evolution (Bazin, et al. 2006; Galtier, et al. 2009; Dong, et al. 2021; Dowling and Wolff 2023). Indeed, mtDNA evolution in arthropods and especially insects, can be significantly impacted by cytoplasmic incompatibilities (CI’s) with endosymbionts like the *Wolbachia* bacteria (Hurst and Jiggins 2005; Galtier, et al. 2009; Dong, et al. 2021; Dowling and Wolff 2023). Furthermore, epistatic interactions between mitochondrial and nuclear genome are also suspected to modulate mtDNA genetic variation given the key biological processes happening in the mitochondria and the tight coordination between the two genome compartments (Wolff, et al. 2014; Sloan, et al. 2015; Rand, et al. 2018; Dowling and Wolff 2023; Nguyen, et al. 2023). For example, an increasing number of studies in mosquitoes suggests that mitochondrial respiration and the associated production of reactive oxygen species (ROS) play a significant role in mosquito immune response and metabolic processes involved in pathogens and insecticides resistance (Van Leeuwen, et al. 2008; Ding, et al. 2020). These epistatic interactions are often neglected in ecology and evolution due to the limited number of studies with adequate datasets to test for these effects. The determinants of mtDNA variations in mosquitoes can thus be multifarious (Hurst and Jiggins 2005; Bazin, et al. 2006; Galtier, et al. 2009; Cameron 2014; Wolff, et al. 2014). Therefore, in many cases, mtDNA does not follow a simple neutral genetic evolution and its use in molecular ecology, metabarcoding, and phylogeographic studies requires a clear assessment of the various factors potentially influencing its evolution. However, doing so necessitate investigating mtDNA variation in combination with nuclear genomic data. This is now increasingly possible thanks to democratization of whole genome re-sequencing and large-scale genomic consortium projects.

The genomic resources provided by two consortia – the MalariaGEN *Anopheles gambiae 1000 genome (Ag1000G)* consortium (The Ag1000G Consortium 2017, 2020) and the *Anopheles* 16 genomes project (Neafsey, et al. 2013; Fontaine, et al. 2015; Neafsey, et al. 2015) – offer a unique opportunity to explore the determinants of mtDNA variation in two sister mosquito species within the *Anopheles gambiae* species complex (AGC): *Anopheles gambiae* and *Anopheles coluzzii*. The AGC is a medically important group of at least 9 closely related and morphologically indistinguishable mosquito sibling species (White, et al. 2011; Coetzee, et al. 2013; Barrón, et al. 2019; Loughlin 2020; Tennessen, et al. 2021). Three members of this African mosquito species complex (*An. gambiae*, *An. coluzzii*, and *An. arabiensis*) are among the most significant malaria vectors in the world, responsible for the majority of the 619,000 malaria-related deaths in 2021, 96% of which occurred in sub-Saharan Africa and impacted primarily children under the age of five (World Health Organization 2022). The ecological plasticity of these three species contributes greatly to their status as major human malaria vectors (Coluzzi, et al. 2002). In contrast to the other AGC species (*An. quadriannulatus, An. merus, An. melas, An. bwambae, An. amharicus, An. fontenllei*) with more confined geographic distributions, these three species have wide overlapping distributions across diverse biomes of tropical Africa. This ecological plasticity in the AGC is attributed to a large adaptive potential, stemming mainly from three major genomic properties: (1) a strikingly high number of paracentric chromosomal inversion polymorphisms segregating in their genome, which are implicated in adaptation to seasonal and spatial environmental heterogeneities related both to climatic variables and anthropogenic alterations of the landscape, in phenotypic variation such as adaptation to desiccation, or even resistance to pathogens like *Plasmodium sp.* or insecticides (e.g., Coluzzi, et al. 2002; Costantini, et al. 2009; Simard, et al. 2009; Cheng, et al. 2012; Ayala, et al. 2017; Riehle, et al. 2017; Cheng, et al. 2018); (2) an exceptional level of genetic diversity identified in natural populations provides a rich material onto which natural selection can act (The Ag1000G Consortium 2017, 2020); and (3) a high propensity for hybridization and interspecific gene flow connecting directly or indirectly the gene pools from all the species over the evolutionary time-scale of the complex (Crawford, et al. 2015; Fontaine, et al. 2015; Thawornwattana, et al. 2018; Müller, et al. 2021).

The species of the AGC radiated within the past 400 to 500 kyrs (Thawornwattana, et al. 2018; Müller, et al. 2021), and the species barriers are not yet fully formed. Although all members of the AGC can be crossed in the laboratory and produce fertile female hybrids but sterile male hybrids (except for *An. gambiae* and *An. coluzzii*), interspecific hybrization rate in nature is supposed to be extremely low (<0.02%) (Pombi, et al. 2017). An exception is the two-sister species *An. gambiae* and *An. coluzzii* which diverged more recently (*ca.* 60 kyr. before present) according to the most recent estimates (Thawornwattana, et al. 2018; Müller, et al. 2021). These two sister species are at an earlier stage of speciation with no post-zygotic isolation detected (reviewed in Pombi, et al. 2017). Hybrid offspring of both sexes are viable and fertile in the laboratory, but strong pre-zygotic and pre-mating isolation barriers have been identified in nature. Hybridization rate is low (*ca.* 1%) across their overlapping distribution range in West Africa, even if high hybridization rates (up to 40%) were reported in the populations from the African “*far west*” (*i.e.* the coastal fringe of Guinea Bissau and Senegambia (estuary of the river Gambia and Casamance in Senegal) (Lee, et al. 2013; Nwakanma, et al. 2013; Pombi, et al. 2017; Vicente, et al. 2017). New unpublished evidence suggests however that these hybrid populations from the African “*far-west”* could be a distinct cryptic hybrid taxon in which diagnostic alleles typically used to discriminate between *An. gambiae* and *An. coluzzii* are still segregating (A. Miles, *pers. comm.*). Nevertheless, despite the low occurrence of contemporary hybridization rates between the members of the AGC, the large geographic overlap in species distributions, together with porous reproductive barriers have resulted in extensive levels for interspecies hybridization over the evolutionary time scales (Fontaine, et al. 2015).

Extensive introgression rates between species of the AGC combined with elevated levels of incomplete lineage sorting (ILS) due to large effective population sizes contributed to maintain high levels of shared polymorphisms and highly discordant phylogenies along the nuclear genome, greatly hampering the identification of the species evolutionary history (Crawford, et al. 2015; Fontaine, et al. 2015; Thawornwattana, et al. 2018; Müller, et al. 2021). Genome-scale studies depicted a highly reticulated evolutionary history of the AGC with outstanding levels of geneflow being detected between the two-sister species – *An. gambiae* and *An. coluzzii,* and also between *An. arabiensis* and the ancestor of *An. gambiae* and *An. coluzzii*. Most of the genomic regions resistant to introgression on their nuclear genome, and thus informative on the species branching order, were mostly identified in portions of the X chromosome and scattered across less than ∼2% of the autosomes (Crawford, et al. 2015; Fontaine, et al. 2015; Thawornwattana, et al. 2018). The lack of any obvious phylogenetic patterns of species structuration at the mitochondrial genome further supported the extensive level of introgression between these three species (Caccone, et al. 1996; Besansky, et al. 1997; Fontaine, et al. 2015; Hanemaaijer, et al. 2018). Additional interspecific introgression signals were also detected between *An. merus* and *An. quadriannulatus*, between *An. gambiae* and *An. bwambae*, and also along the ancestral branches of the AGC species (Thelwell, et al. 2000; Crawford, et al. 2015; Fontaine, et al. 2015; Thawornwattana, et al. 2018; Müller, et al. 2021). Although the selective and evolutionary effects associated with this extensive level of introgression between species of the AGC remains to be fully investigated, clear evidence of adaptive introgression were detected involving chromosomal inversions (Fontaine, et al. 2015; Riehle, et al. 2017; Thawornwattana, et al. 2018) and insecticide resistance loci (Clarkson, et al. 2014; Grau-Bové, et al. 2020; Grau-Bové, et al. 2021; Lucas, et al. 2023).

Here we leveraged the genomic resources from The Ag1000G Consortium (2017, 2020) and from the *Anopheles* 16 genomes project (Neafsey, et al. 2013; Fontaine, et al. 2015; Neafsey, et al. 2015) to explore the determinants of mitochondrial genetic variation in the *An. gambiae complex* (AGC) with a particular focus on *An. gambiae* and *An. coluzzii*. For that purpose, we first assembled mitogenomes for 1219 pan-African mosquitoes (Fig. 1 and S1) using a new flexible bioinformatic pipeline, called *AutoMitoG* [*automatic mitogenome assembly*] (Fig. S2), which relies on *MitoBIM* approach that combines mapping and *de-novo* assembly of short-read sequencing data (Hahn, et al. 2013). We then assessed the level of mtDNA variation, its phylogeographic and population structure, and the mtDNA genealogical history in comparison with the population demographic history previously estimated from the nuclear genome (The Ag1000G Consortium 2017, 2020). We further assessed what factors best explain the mtDNA phylogeographic structure, testing various covariates including *Wolbachia* infection status, population structure estimated from nuclear genome data, and chromosomal inversions. Finally, we investigated the mito-nuclear associations that possibly imply coevolution and coadaptation between the two genomic compartments.

**Figure 1.**
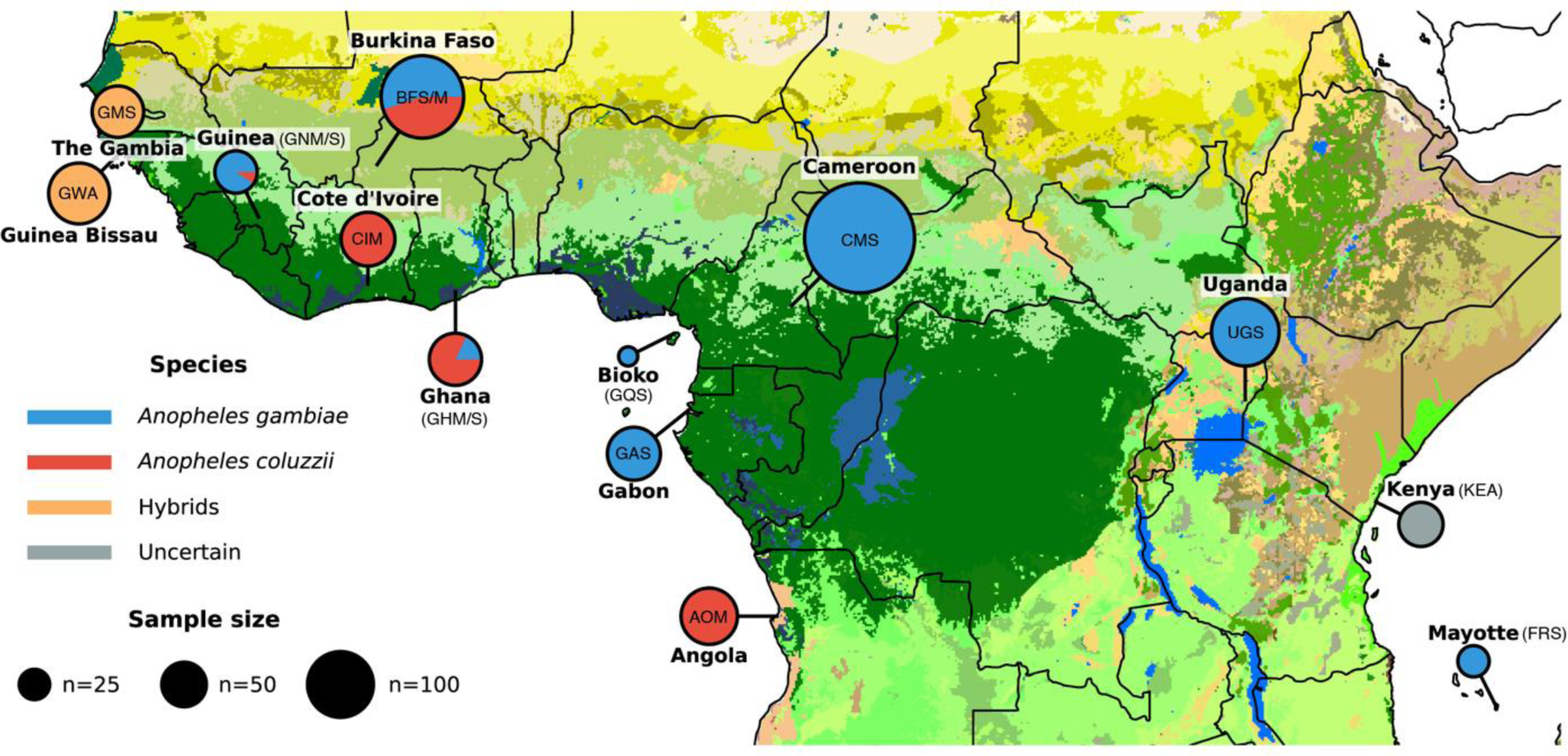
Approximate sampling locations and sample size per location of the 1142 samples of *An. gambiae* and *An. coluzzii* from the The Ag1000G Consortium (2020). The population codes are also provided within or next to the pie-charts (see Table 1 and S2). Colors within the pie-charts describe the species and include: An. *gambiae* (in blue; formerly the *S-form* of *An. gambiae*), *An. coluzzii* (in red; formerly known as the M-form of *An. gambiae*), the hybrid taxonomically uncertain populations of the African *far-west* (in orange), and the taxonomically uncertain population of Kenya (in grey). The figure is modified from The Ag1000G Consortium (2020). Map colors represent ecosystem classes; dark green designates forest ecosystems. For a compete color legend see Figure 9 in the work of Sayre (2013). (see Table S2 for further details on the sampling).

## Results and discussion

### The *AutoMitoG* pipeline and assembly of the Ag1000G mitogenomes

The *AutoMitoG* pipeline, which streamlines mitogenome assembly using the MitoBIM approach (see the method section and Fig. S2), successfully assembled 1219 mitogenome sequences from the unmapped short read data originating from the two *An. gambiae* consortia projects (Fontaine, et al. 2015; The Ag1000G Consortium 2017, 2020) (Fig. 1 and S1, Table S1 and S2). We first assessed the pipeline performance by comparing newly assembled mitogenome sequences with those from the 74 samples of the AGC previously generated in Fontaine, et al. (2015) (Fig. S1, Table S1). Average assembly length before any trimming and sequence alignment was 15,366 base pairs (bp). Following Fontaine, et al. (2015) and after sequence alignment, we removed the control region (CR) resulting in a 14,843 bp alignment length. Excluding the CR removed most of the ambiguities and gaps remaining in the alignment (Fig. S3). The previous and present bioinformatic pipelines generated very similar mtDNA assemblies for each sample with one exception (samples ID: Aara_SRS408148, Fig. S4). Beside that sample which resulted from a label mistake in the DRYAD repository of Fontaine, et al. (2015), mitogenome sequence pairs for each sample were nearly identical with a number of nucleotide differences of 0.4 on average (25% quartile: 0.0; 75% quartile: 1.0, max: 3.0) (Fig. S4). We augmented this alignment with newly assembled mitogenome sequences from three *An. bwambae* samples of lower sequencing quality than the other samples (Table S1). These mitogenomes assembled with the two pipelines also generated similar sequences with a slightly lower sequence identity (>99.9%) for each pair of assemblies, except one sample (bwambae_4) which was more difficult to assemble (Fig. S4a and Table S1). Overall, the *AutoMitoG* pipeline performed at a good bench mark level.

We then applied the *AutoMitoG* pipeline to the 1142 *An. gambiae* and *An. coluzzii* mosquito samples of The Ag1000G Consortium (2020)) (Fig.1, Table S2). Assembly lengths were 15,364 ± 1.9 bp on average (min: 15,358 – max: 15,374) (Table S2). The raw alignment was 15,866 bp long, and 14,844 bps after removing the CR and gaps. The 1142 mtDNA sequence alignment of *An. gambiae* and *An. coluzzii* included 3017 polymorphic sites (S), 1195 singleton sites (*Sing.*), and a nucleotide diversity (π) of 0.004, defining 910 distinct haplotypes (H), with a haplotype diversity (HD) of 0.999 (Table 1).

**Table 1.** Mitochondrial genetic diversity statistics per species and population for the entire mitogenome alignment (14,844 bps). Number of sequences (N), INDEL sites (INDELS), segregating sites (S), singletons (*Sing.*), shared polymorphism (*Shared P.*), average number of differences between pairs of sequences (K), number of haplotypes (H), haplotype diversity (HD), nucleotide diversity (π), Theta-Waterson (Θ _W_), Tajima’s D, number (and proportion) of sequences belonging to the cryptic lineage (N (%) Cryptic).

| Species | Group | Location | N | INDELS | S | Sing. | Shared P. | K | H | HD | $\pi$ | $\Theta_w$ | Tajima's D* | N (%) Cryptic |
| --- | --- | --- | --- | --- | --- | --- | --- | --- | --- | --- | --- | --- | --- | --- |
| All | All | All | 1142 | 14 | 3017 | 1195 | 1822 | 57.5 | 910 | 0.999 | 0.0039 | 0.0266 | -2.50* | 232 (20%) |
| Hybrid | Hybrid | Hybrid | 156 | 2 | 942 | 468 | 474 | 53.8 | 131 | 0.997 | 0.0036 | 0.0113 | -2.23* | 77 (49%) |
| *An.coluzzii* | *An.coluzzii* | *An.coluzzii* | 283 | 4 | 1298 | 541 | 757 | 60 | 237 | 0.998 | 0.0040 | 0.014 | -2.24* | 89 (31%) |
| *An.gambiae* | *An.gambiae* | *An.gambiae* | 655 | 10 | 2336 | 1004 | 1332 | 54.3 | 539 | 0.999 | 0.0037 | 0.0222 | -2.51* | 66 (10%) |
| *An.coluzzii* | AOM | Angola | 78 | 2 | 323 | 121 | 202 | 37.5 | 50 | 0.983 | 0.0025 | 0.0044 | -1.48 | 4 (5%) |
| *An.coluzzii* | BFM | Burkina Faso | 75 | 1 | 819 | 489 | 330 | 59.7 | 75 | 1.000 | 0.0040 | 0.0113 | -2.25* | 37 (49%) |
| *An.coluzzii* | CIM | Côte d'Ivoire | 71 | 0 | 552 | 260 | 292 | 58.6 | 58 | 0.988 | 0.0040 | 0.0077 | -1.71* | 27 (38%) |
| *An.coluzzii* | GHM | Ghana | 55 | 1 | 498 | 224 | 274 | 61.6 | 53 | 0.998 | 0.0041 | 0.0073 | -1.57* | 20 (36%) |
| *An.coluzzii* | GNM | Guinea | 4 | 0 | 61 | 61 | 0 | 30.5 | 2 | 0.500 | 0.0021 | 0.0022 | -0.87* | 1 (25%) |
| *An.gambiae* | BFS | Burkina Faso | 92 | 2 | 1019 | 585 | 434 | 56.9 | 91 | 1.000 | 0.0038 | 0.0135 | -2.46* | 10 (11%) |
| *An.gambiae* | CMS | Cameroon | 297 | 8 | 1537 | 639 | 898 | 56.6 | 217 | 0.996 | 0.0038 | 0.0164 | -2.41* | 47 (16%) |
| *An.gambiae* | FRS | Mayotte | 24 | 0 | 31 | 18 | 13 | 5.3 | 15 | 0.920 | 0.0004 | 0.0006 | -1.35 | 0 (0%) |
| *An.gambiae* | GAS | Gabon | 69 | 0 | 283 | 117 | 166 | 38.3 | 48 | 0.972 | 0.0026 | 0.004 | -1.23 | 2 (3%) |
| *An.gambiae* | GHS | Ghana | 12 | 0 | 215 | 149 | 66 | 48.4 | 11 | 0.985 | 0.0033 | 0.0048 | -1.5 | 1 (8%) |
| *An.gambiae* | GNS | Guinea | 40 | 2 | 574 | 357 | 217 | 57.3 | 38 | 0.997 | 0.0039 | 0.0091 | -2.16* | 4 (10%) |
| *An.gambiae* | GQS | Bioko Island | 9 | 0 | 78 | 48 | 30 | 22.8 | 9 | 1.000 | 0.0015 | 0.0019 | -1.06 | 0 (0%) |
| *An.gambiae* | UGS | Uganda | 112 | 3 | 949 | 519 | 430 | 49.8 | 112 | 1.000 | 0.0034 | 0.0121 | -2.44* | 2 (2%) |
| Hybrid | GMS | The Gambia | 65 | 0 | 364 | 140 | 224 | 47.8 | 43 | 0.982 | 0.0032 | 0.0052 | -1.33 | 42 (65%) |
| Hybrid | GWA | Guinea Bissau | 91 | 2 | 792 | 436 | 356 | 55.7 | 88 | 0.999 | 0.0038 | 0.0105 | -2.20* | 35 (38%) |
| Uncertain | KEA | Kenya | 48 | 0 | 81 | 1 | 80 | 32.3 | 4 | 0.650 | 0.0022 | 0.0012 | 2.74 | 0 (0%) |

\* Star on Tajima's D values indicates a significant departure from neutrality ( $D \neq 0$ ) based on 1000 coalescent simulations.

### Phylogenetic relationships among mtDNA haplotypes trace the phylogeographic history of the species split and introgression among species of the *An. gambiae* complex

Phylogenetic relationships among the 1142 mtDNA sequences from *An. gambiae* and *An. coluzzii*, together with the 77 mtDNA sequences from the five other species from the AGC (Figure 2, see also Fig. S4 and S5) were consistent with previous studies. Indeed, previously reported evidence of extensive gene flow between *An. gambiae*, *An. coluzzii*, and *An. arabiensis* found support in our phylogenetic analyses with a complete absence of any mtDNA haplotype private to *An. arabiensis* samples (Figure 2, see also Fig. S4 and S5). From the mtDNA standpoint, the 12 samples of *An. arabiensis* could not be discriminated from *An. gambiae* and *An. coluzzii* as previously reported (Besansky, et al. 1997; Donnelly, et al. 2001; Fontaine, et al. 2015; Hanemaaijer, et al. 2018). Aside from *An. gambiae*, *An. coluzzii*, and *An. arabiensis*, the four other species of the AGC clustered in a monophyletic divergent clade. Within that clade, the three salt-water tolerant species (*An. melas*, *An. merus*, and *An. bwambae*) formed a monophyletic group next to *An. quadriannulatus*. One *An. bwambae* sample (bwambae3, Fig. 2 and S4) carried a mtDNA haplotype clustering among those from *An. gambiae*, *An. coluzzii* and *An. arabiensis.* This is consistent with previous evidence of mitochondrial introgression between *An. bwambae* and one of these three species, most likely *An. gambiae* (Thelwell, et al. 2000). Noteworthy, one *An. gambiae* specimen from Cameroon (AN0293_C_CMS, Fig. 2A) carried a unique mitogenome haplotype closely related to *An. quadriannulatus*. This peculiar haplotype likely reflects the historical/original *An. arabiensis* mitogenomes before being fully replaced by those of *An. gambiae* and/or *An. coluzzii* and still segregating at low frequency in the gene pool of the three species. This finding of a “relict” haplotype from the non-recombining mtDNA locus is consistent with the admitted species branching order of the *An. gambiae* species complex, as depicted by the X chromosome (Fontaine, et al. 2015; Thawornwattana, et al. 2018; Müller, et al. 2021). The phase-3 of The Ag1000G Consortium (2021), which includes hundreds of samples from *An. arabiensis*, will provide further insights on this topic.

**Figure 2.**
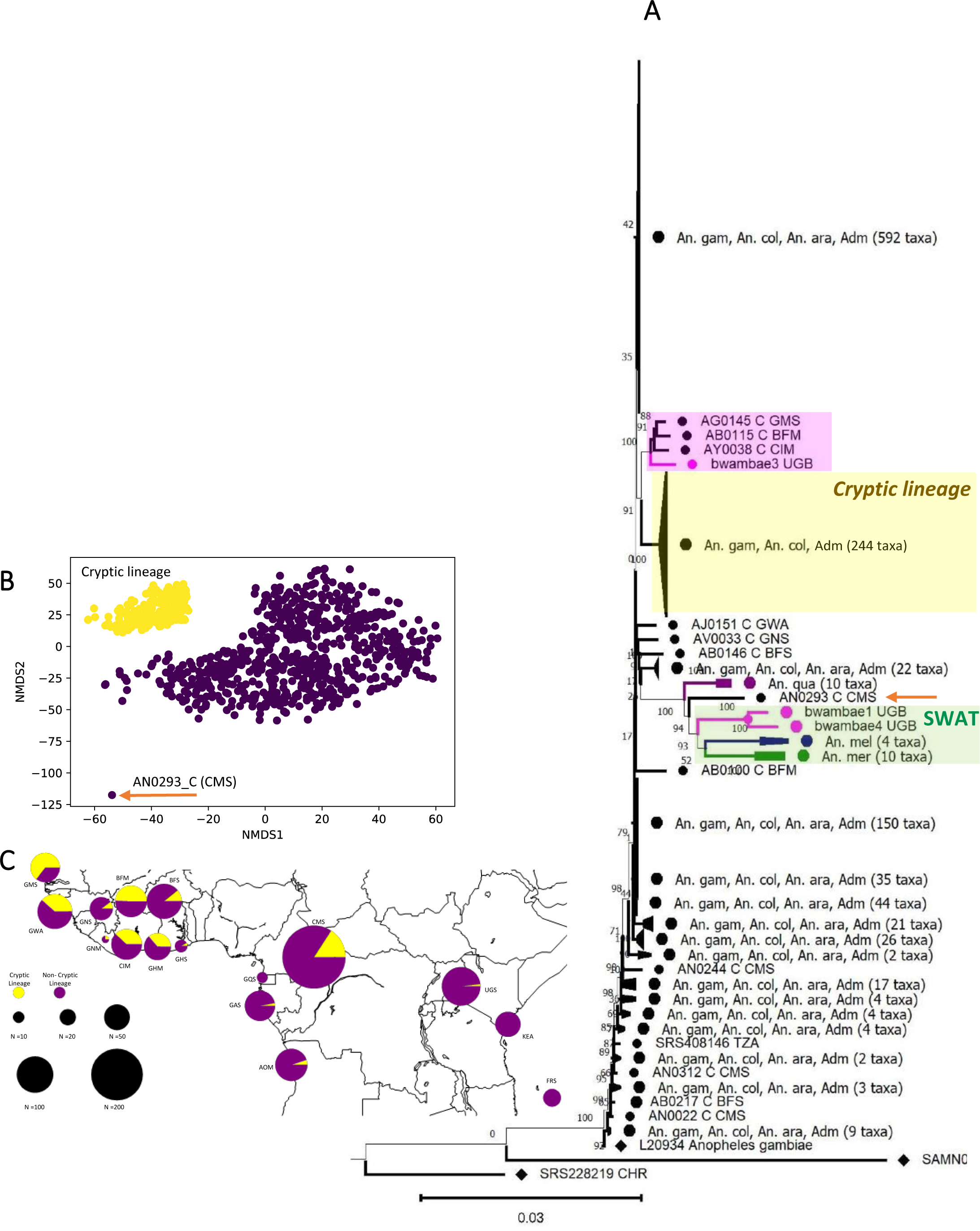
Phylogenetic relationships among mitogenomes. (A) MtDNA maximum likelihood phylogeny based on the 1222 sequences composed of the 1142 *An. gambiae and An. coluzzii* samples from the Ag1000G, 77 from the 7 species of the AGC from Fontaine et al (2015), on reference sequence from *An. gambiae*, and the two outgroups. (Black dots: An. *gambiae (An. gam), An. coluzzii (An. col), or An. arabiensis (An. ara), Green: An. merus (An. mer), Blue: An. melas (An. mel), Purple, An. quadriannulatus (An. qua), Pink: An. bwambae).* Bootstrap support values (%) are indicated at the nodes. Notice the absence of *An. arabiensis* specific mtDNA haplotype, the monophyletic clustering of the salt-water tolerent species (SWAT in green), the introgressed bwambae3 sample within within *An. gambiae* and *An. coluzzii* clade (in pink), and the presence of one *An. gambiae* individual (AN0293 CMS, orange arrow) next to *An. quadriannulatus,* in a place where *An. arabiensis* would be expected based on the admitted species tree (Fontaine et al. 2015). (B) Distance-based nMDS analysis of the 1142 mtDNA sequences from the Ag1000G with a color-coding based on a hierarchical clustering analysis. The cryptic lineage is in yellow. The outlier sample at the bottom is AN0293_C CMS is most likely a relic of *An. arabiensis* haplotype (see Fig. 2A). None of the samples associated with bwambae3 (AB0015 from BFM, AG0145 from GMS, and AY0038 CIM) were found in the cryptic lineage. (C) Geographic distribution of the cryptic lineage.

While most of the mtDNA haplotypes carried by *An. gambiae*, *An. coluzzii*, and *An. arabiensis* were closely related, as shown by the short branches on the phylogenetic tree (Fig. 2A, Fig. S4 and S5) and on the distance-based non-metric multidimensional scaling (nMDS) (Fig. 2B, Fig. S6), a group of 244 samples (232 from the The Ag1000G Consortium (2020) and 12 from Fontaine, et al. (2015)) clustered into a distinctive clade (hereafter called the “cryptic lineage”) (Fig. 2A and 2B). This cryptic lineage displayed a higher level of divergence than the others within the mtDNA gene pool of *An. gambiae, An. coluzzii*, *An. arabiensis*, yet similar-to-slightly-lower than for the clades containing the other 4 species of the AGC (Fig. 2A). Interestingly, the geographic distribution of this cryptic group matched closely with the geographic distribution of *An. coluzzii*, mostly prevalent in the African “*far-west*” side of the distribution of the two species, and decreasing in frequency eastwards and southwards (Fig. 2C). The prevalence of the cryptic lineage was the most important in the hybrids (taxonomically uncertain) populations where it reached *ca.* 50% of the samples (up to 65% for the populations of the Gambia (GMS) and 40% of the Guinea-Bissau (GWA)), then composing 31% of the *An. coluzzii* samples, and less than 10% of the *An. gambiae* samples (Table 1, Fig. 2C, Fig. S6). This clear enrichment of the cryptic mtDNA lineage in the populations from the African *far-west*, especially in the hybrid and *An. coluzzii* populations, its level of divergence compared to the common mtDNA lineage which was comparable to the levels observed among species of the AGC (Fig. 2), together with the West to East gradient decline, all these observations suggest it could be a phylogeographic legacy of the split between the two sister species, followed by a secondary contact with an incomplete homogenization of the mtDNA gene pool.

### Significant mtDNA genetic structure among and within *An. gambiae* and *An. coluzzii*

An Analysis of Molecular Variance (AMOVA) (Excoffier, et al. 1992) showed that most of the mtDNA variation was distributed within populations (87.0%), but significant variance partitioning was also observed between populations (9.6%, p<0.001), and between species as well (3.4%, p<0.007) (Table 2). Level of population differentiation (Fig. S7), expressed as *F_ST_*values among populations between nuclear genome (nuDNA) from The Ag1000G Consortium (2020) and mtDNA genome were strongly correlated (Fig. S8). *F_ST_*values at the nuclear genome explained *ca.* 80% of the mtDNA *F_ST_*values (*p* < 0.001). All comparisons involving the isolated island population of Mayotte (FRS) displayed both high values at the nuDNA and mtDNA genome, the highest mtDNA values being observed between the island populations of Mayotte (FRS) and Bioko (GQS) (Fig. S7 and S8). Such elevated levels of mtDNA and nuDNA differentiation reflect the small long-term effective population size and limited gene flow, with potential repeated bottleneck/founder effects. All these contribute to a strong genetic drift of this Mayotte Island populations (FRS), as previously reported (The Ag1000G Consortium 2020). Globally, genetic differentiation observed at the mtDNA were overall higher than those at the nuclear genome, which likely reflect the reduced effective size of the mtDNA compared to the nuDNA. Exceptions included all comparisons involving the taxonomically uncertain and very peculiar population of Kenya (KEA), where the *F_ST_* values were lower or equivalent (Fig. S7 and S8). Overall, we observed a high concordance between the mtDNA and nuDNA levels of population differentiation. These results further underline that both geographical location and, to a lesser extent, species differentiations within and between *An. gambiae* and *An. coluzzii* are major determinants of mtDNA variation.

**Table 2.**
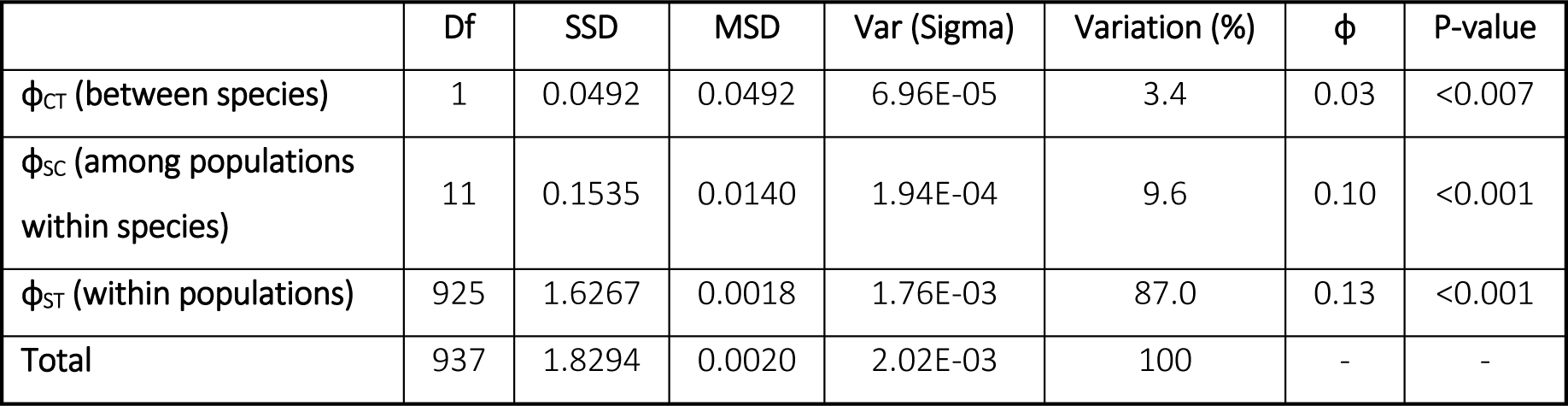
Analysis of Molecular Variance (AMOVA) describing the variance partitioning at three hierarchical levels: between species, between populations within species, and within populations. The AMOVA was conducted with the TN93+gamma model of sequence evolution. The analysis was conducted considering only populations from the Ag1000G that were taxonomically unambiguous (n=938, see Fig. 1), thus removing the uncertain populations from The Gambiae (GMS) and Guinea-Bissau (GWA), as well as the population from Kenya (KEA). The table provide the main results of the AMOVA including the degree of freedom (Df) at each level, the sum square and mean square deviations (SSD and MSD), variance component (*σ*), and variance proportion (%), ɸ-statistics, and *p-*value from 1000 permutation test.

### MtDNA isolation-by-distance patterns reflect distinct life histories between *An. gambiae* and *An. coluzzii*

Previous studies showed that genetic differentiation (i.e., *F_ST_* or its linearized equivalent *F_ST_*/(1-*F_ST_*)) at the nuclear genome significantly increased with geographic distance in *An. gambiae* and *An. coluzzii* (Lehmann, et al. 2003; The Ag1000G Consortium 2020). This isolation-by-distance (IBD) pattern was significantly stronger in *An. coluzzii* than in *An. gambiae*, translating into reduced local effective population size and/or reduced intergenerational dispersal distance in the first compared to the second species (see Fig. 3 in The Ag1000G Consortium 2020). In line with these findings at the nuclear genome, we found significant IBD at the mtDNA as well when considering all populations irrespective of the species (Mantel’s *r =* 0.35; *p* < 0.003; n=13), with a very strong signal among populations of *An. coluzzii* (Mantel’s *r =* 0.96; *p* = 0.017; n=5), and a weaker marginal signal among populations of *An. gambiae* (Mantel’s *r =* 0.32; *p* = 0.095; n=8) (Table S4). However, these analyses included populations that were found genetically isolated by geographic barrier to geneflow when analyzing the nuclear genome (Angola – AOM in *An. coluzzii*; Gabon - GAS and Mayotte Island - FRS in *An. gambiae*) (The Ag1000G Consortium 2020). These geographic barriers to dispersal can artificially inflate the IBD patterns without necessarily implying reduced neighborhood size, which is the product of reduced local effective population density and intergenerational dispersal distance that increase local genetic drift (Wright 1946; Rousset 1997). The IBD signal among populations within species becomes weaker and not statistically different from zero when removing geographically isolated populations (AOM, GAS, or FRS) from the IBD analysis (*An. coluzzii* Mantel’s *r*=0.46, p=0.167, n=4; *An. gambiae* Mantle’s r=-0.18; p=0.617; n=6). Nevertheless, despite the lack of significant results likely due to the small number of sampled populations, the strength of association between genetic and geographic distances still remain strong and positive in *An. coluzzii* with a *r^2^* value of 0.21, which is very comparable to the *r^2^* value of 0.22 observed at the nuclear genome (see Fig. 3B in The Ag1000G Consortium 2020). This contrasts with the lack of any detectable IBD signal at the mtDNA genome among populations of *An. gambiae* and the very weak IBD signal found on the nuclear genome. These results are consistent with the distinct life history and dispersal strategies between the two species (Dao, et al. 2014; Huestis, et al. 2019; Hemming-Schroeder, et al. 2020; Faiman, et al. 2022). A significant fraction of the populations of *An. coluzzii* from NW Africa seem to endure locally the dry season by engaging into aestivation strategy to rebound from local founders when the wet season starts. In contrast, *An. gambiae* populations go locally extinct during the dry season and rebound after a certain lag time by long-distance migration. Since female mosquitoes potentially disperse more and live also longer than males (Yaro, et al. 2022), we may have expected weaker evidence of IBD at the mtDNA compared to the signal found at the nuclear genome. However, we did not observe this effect. Thus, if this effect exists, it would likely be counter balanced by the strong differences in aestivation and dispersal strategies between the two species.

**Figure 3.**
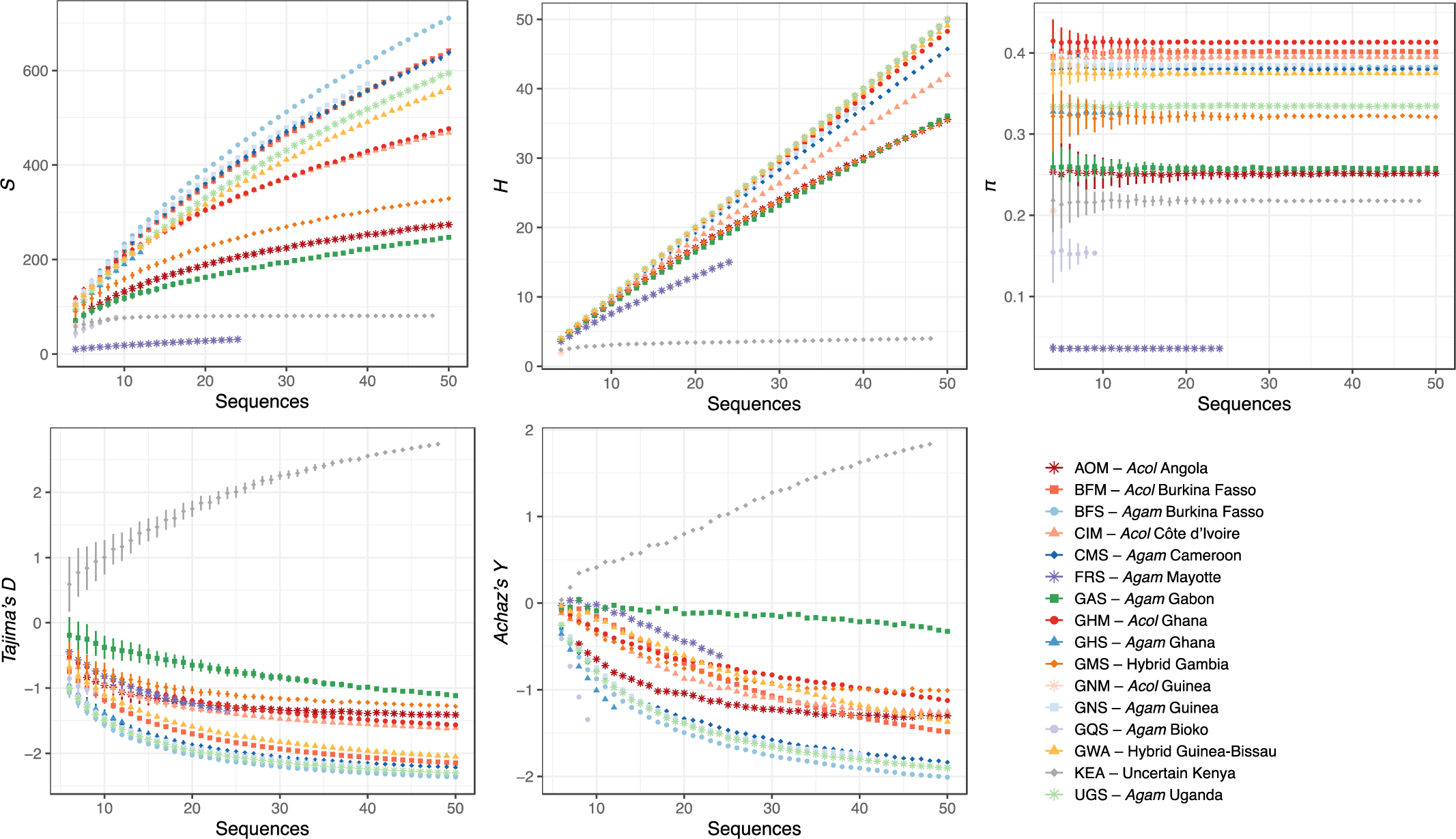
Mitochondrial genetic diversity statistics for each population of the Ag1000G. The statistics shown include the number of segregating site (*S*), the number of haplotypes (*H*), the nucleotide diversity (*π*), the Tajima’s *D*, and the Achaz’s *Y*. The rarefaction curves describe the impact of varying sample size on the estimated values for each statistic and for each population. The mean and standard error values are reported for each sample size increment from 3 to 50.

### MtDNA variation in line with population demography of *An. gambiae* and *An. coluzzii*, but with an imprint of the cryptic lineage history

Patterns of mtDNA variation among populations of *An. gambiae* and *An. coluzzii* (Table 1, Fig. 3) were consistent with those previously reported at the nuclear genome (The Ag1000G Consortium 2020). The exceptional genetic diversity previously observed at the nuclear genome also manifested at the mtDNA level by a high overall level of haplotype diversity (*HD*=0.999), with 910 distinct haplotypes found in 1142 samples, an average number (K) of 58 differences between pairs of haplotypes, and 3017 segregating sites including one third of singletons (Table 1).

Rarefaction curves, which account for differences in population sample sizes, for the number of segregating sites (*S*) and the number of haplotypes (*H*) kept increasing with the sample size in most populations. These curves clearly showed that the plateau was not within reach with a sampling up to 50, especially for the populations located North of the Congo River Basin and West to the Rift Valley (Fig. 3). In term of nucleotide diversity (π), these populations were also among the most diversified, with the highest values observed for the *An. coluzzii* populations from the NW Africa, followed by the *An. gambiae* populations from the same regions, and the hybrid (taxonomically uncertain) population in the Guinea Bissau (GWA). These high levels of nucleotide diversity (π) actually reflected populations in which there was a mixed proportion of haplotype from the cryptic and common mito-groups identified in the phylogenetic analyses (Fig. 2). Nucleotide diversity (π) decreased in populations where the haplotype mixture between cryptic and common lineages decreases, for example in the hybrid (taxonomically uncertain) population of Gambia (GMS) where the cryptic lineage dominates or in the *An. gambiae* population of Uganda (UGS) where it is almost absent. Overall, the high level of mtDNA variation combined with very negative values for Tajima’s D or Achaz’s Y statistic (Achaz 2008) indicate an excess of rare variants. These results support previous demographic inference modelling, showing large effective population sizes in the NW Africa distribution ranges of the two species and evidence for historical population expansions (The Ag1000G Consortium 2017, 2020). These conditions where genetic drift is very ineffective are highly favorable to maintain high genetic diversity.

The (semi-)isolated populations from Gabon (*An. gambiae –* GAS) and Angola (*An. coluzzii* – AOM) displayed intermediate values of genetic diversity and Tajima’s D and Achaz’s Y values closer to zero (Table 1, Fig. 3). This is consistent with a historically more stable population size, and reduced effective size as previously reported (The Ag1000G Consortium 2017, 2020). The two *An. gambiae* populations from the islands of Mayotte (FRS) and Bioko (GQS) also departed from the other populations at the mtDNA variation with very low nucleotide and haplotype diversity, and slightly negative Tajima’s D and Achaz’s Y values. These are further evidence for small effective population size, and suggestive of strong bottlenecks (or founder effects). In these island populations, the number of haplotypes was small and closely related to each other, with excess of rare variants, as expected after strong bottlenecks which can result from cyclic variation in population sizes, with possibly repeated founder events.

The taxonomically uncertain population from Kenyan (KEA) was already known for its very peculiar patterns of genetic diversity at the nuclear genome, with specificities close to a colony population with mixed ancestry from *An. gambiae* and *An. coluzzii* (see Fig. 4 in The Ag1000G Consortium 2020). The Kenyan population was also an outlier population at the mtDNA genome with only four distinct haplotypes detected that differ from each other at only ∼32 sites with almost no singletons, thus a very low haplotype diversity (HD=0.65) compared to the other populations, and the only population in the Ag1000G sampling with highly positive Tajima’s *D* and Achaz’s *Y* values (Table 1, Fig. 3).

**Figure 4.**
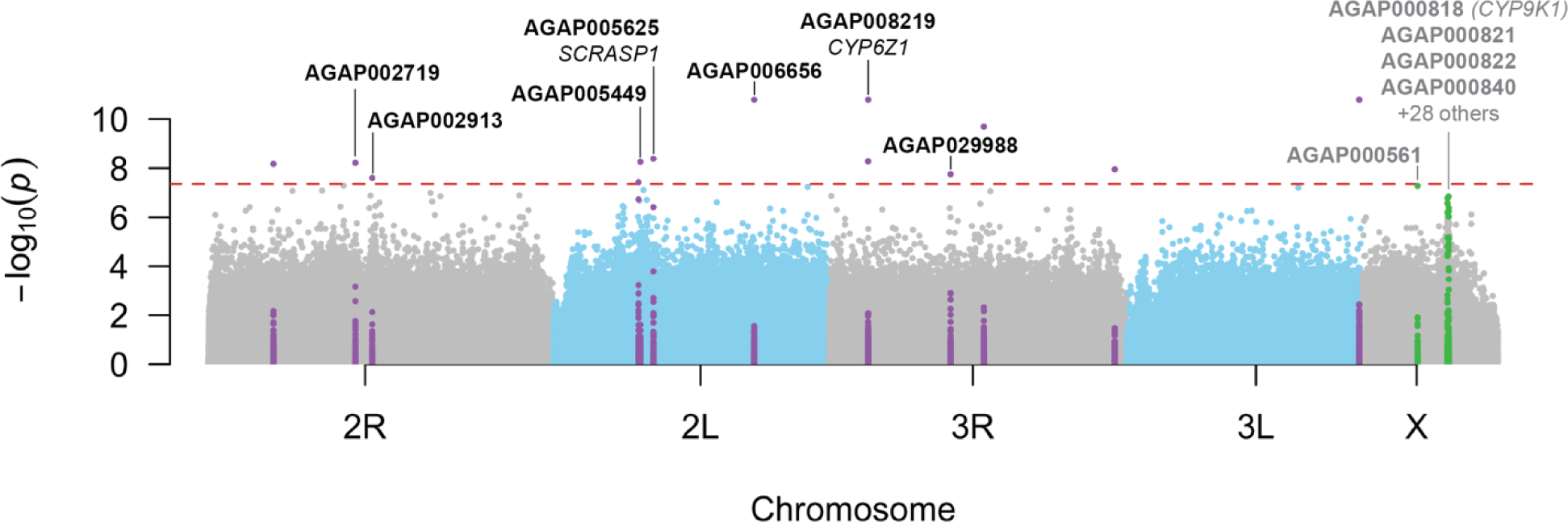
Manhattan plot showing the genetic associations between the two major mtDNA lineages and each of the SNPs on the nuclear genome. The GWAS was conducted accounting for population structure using the 6 first PCs, sex, and *Wolbachia* occurrence as covariates. Each chromosome arms are colored-coded. The red dash horizontal line shows the Bonferroni corrected significance threshold of 4.4 10^-8^. The 14 significant SNPs together with the SNPs 1kb upstream or downstream are marked in purple. SNPs in green are those marginally significant on the X chromosome forming a clear “skyscraper”. Gene-ID and gene name, when an annotated transcript was available, are displayed. See fig. S14 for QQ-plot, fig. S15 for a zoomed view of each significant SNPs, and fig. S16 for a zoomed view on the X-chromosome.

### The cryptic mtDNA lineage: a phylogeographic legacy of the split between *An. gambiae* and *An. coluzzii*

We investigated further the specificities of the distinctive cryptic mtDNA lineage (Fig. 2) to better understand its potential evolutionary origin(s). We tested whether its occurrence was associated with the potential occurrence of *Wolbachia* infection, the population genetic structure at the nuclear genome, as estimated using a principal component analyses (PCA) following The Ag1000G Consortium (2017, 2020), and other genomic features previously characterized for these samples, including major chromosomal inversions (2L^a^, 2R^b,c,d,u^), and insecticide resistance mutations (*rdl226*, *vgsc995*).

The intracellular and intraovarian *Wolbachia* bacterium is frequently found in insects and can be a strong manipulator of insect reproductive biology, impacting physiology, behavior, creating cytoplasmic incompatibilities, and could even act as a speciation agent (Rokas 2000; Werren, et al. 2008; Galtier, et al. 2009; Bruzzese, et al. 2021; Dong, et al. 2021; Dowling and Wolff 2023). *Wolbachia* can thus have significant impacts on mitochondrial heritability and its genetic variation. It was previously detected in *An. gambiae* and *An. coluzzii*, even though the vertical transmission or impacts on the reproductive biology of these mosquitoes is still debated (Baldini, et al. 2014; Shaw, et al. 2016; Gomes, et al. 2017; Gomes and Barillas-Mury 2018; Jeffries, et al. 2018; Pascar and Chandler 2018; Ayala, et al. 2019; Chrostek, et al. 2019; Straub, et al. 2020; Bamou, et al. 2021; Jeffries, et al. 2021).

We used the method of Pascar and Chandler (2018) to detect *Wolbachia* occurrence, using the unmapped Illumina short-read data of the The Ag1000G Consortium (2020). Using a lenient set of filters (at least 3 reads mapping to *Wolbachia* sequence with at least 90bp and 90% sequence identity), we found 111 (9.7%) individual mosquitoes carrying reads blasting to the *Wolbachia* supergroup A (Fig. S9 and Table S5). This detection rate dropped to 27 (2.4%) positive individuals when using stricter detection filters (3 reads blasting to Wolbachia sequences with at least 98bp length and 95% identity) similar to those used by Pascar and Chandler (2018). *Wolbachia* was primarily detected in the *An. gambiae* population of Mayotte (FRS; lenient: 83% or strict: 17%), and in the *An. coluzzii* populations of Côte d’Ivoire (CIM; lenient: 55% or strict: 23%) and Ghana (GHM; lenient: 44% or strict: 4%) (Fig. S9 and Table S5). These infection rates were quite low, especially if we consider the stricter criteria of Pascar and Chandler (2018). These rates were in line with previous reports by Chrostek, et al. (2019) who even questioned the natural occurrence of *Wolbachia* in natural populations of *An. gambiae* and *An. coluzzii.* Chrostek, et al. (2019) argued that such a low number of reads could come from ingested food, or mosquito parasites infected by *Wolbachia* (e.g. nematodes). We did not find any significant association between the *Wolbachia* potential occurrence and the cryptic mtDNA lineage (ranked predictive power of cross-features *x2y* metric = 0; Fig. S10).

Population genetic structure was estimated by a PCA on 100k independent single nucleotide polymorphisms (SNPs) from the nuclear genome (Fig. S11), following the same procedure as in The Ag1000G Consortium (2017, 2020). The PC1, PC6 and (to a lesser extent) PC2 were significant predictors of the cryptic mtDNA lineage occurrence, explaining between 13% and 15% of the cryptic mtDNA lineage variation for PC1, 21% for PC6, and 2% for PC2 (Fig. S10). PC1 discriminates *An. coluzzii* from *An. gambiae*, PC6 splits the hybrid (taxonomically uncertain) populations from the other populations of *An. coluzzii* and *An. gambiae*, and PC2 reflects the strong differentiation of the Angolan (AOM) population from the other *An. coluzzii* and *An. gambiae* populations (Fig. S11). Altogether, these associations between the PCs and the cryptic mtDNA lineage occurrence underline its variation according to species and geography visually displayed in Fig. 4. Beside the PCs, no other genomic features tested here significantly correlated with the occurrence of the cryptic mtDNA lineage variation in natural populations, except for the 2La chromosomal inversion frequency. However, the 2La inversion was also strongly associated with the population genetic structure capture by the PCs, suggesting its association with the cryptic mtDNA lineage could be an “echo” of the population genetic structure (Fig. S10).

Taken together, all the above results suggest that the cryptic mtDNA lineage is likely a phylogeographic legacy of the split between *An. colluzzii* and *An. gambiae*. It likely arose during a period of isolation in *An. coluzzii,* as suggested by its level of divergence similar to the interspecific mtDNA divergence observed between species of the AGC, and by the enrichment of this lineage in the populations of *An. coluzzii* and in the taxonomically uncertain populations from the African *far-west* (Fig. 2). Together with the West to East gradient decline, these results suggest that the two sister species went back into contact with an incomplete homogenization of the mtDNA gene pool.

### Mito-nuclear interactions suggest selection on the mito-group divergence related to metabolic resistance to pathogens and insecticides

Selection may have been involved in preventing a full homogenization of the mtDNA gene pool(s) between *An. coluzzii, An. gambiae*, and the hybrid (taxonomically uncertain) populations, with possible mito-nuclear interactions. To test this hypothesis, we conducted a genome-wide association study (GWAS), testing which SNPs on the nuclear genome were significantly associated with the cryptic mtDNA lineage occurrence (which is considered here as a binary variable). The GWAS was conducted considering covariates including the 6 first PCs (Fig. S11) to account for population genetic structure, *Wolbachia* occurrence (Table S5 and Fig. S9), and sex. As such analysis requires unrelated samples (Uffelmann, et al. 2021), we excluded closely related sample pairs in the Ag1000G dataset with kinship coefficient exceeding the level of 2^nd^ degree relatives (Table S6). The KING-robust method (Manichaikul, et al. 2010), which relaxes the assumption of genetic homogeneity within population, identified multiple related pairs of samples within populations equal or exceeding the level of 2^nd^ degree relatives, with some cases of full-sib or parent-offspring’s, and even rare cases of monozygotic twins between pairs of mosquitoes (see Fig. S12, S13, Table S7, and S8). Full-siblings or parent-offspring’s relationships in mosquitoes can occur if samples originated from larvae from a single female for example. Monozygotic twin’s relationship can either reflect sample duplicates in the dataset or highly inbred samples as would be observed in samples coming from a laboratory colony. Unsurprisingly, the most impacted population was the Kenyan (KEA) one. Its peculiar genetic make-up is similar to a laboratory colony, as was previously spotted in The Ag1000G Consortium (2017). However, instances of full siblings and monozygotic twins were found in the populations from Cameroon (CMS) and Angola (AOM) (see Fig. S12, S13, Table S7, and S8). Overall, removing 98 samples from the dataset (Table S9) resolved all the issues allowing only up to the 3^rd^ degree relative association between sample pairs. The cleaned SNPs dataset used in the GWAS included 1044 unrelated samples (Fig. S11b) and 7,858,575 nuclear biallelic SNPs (Table S10). After removing related samples, and accounting for population structure, sex, and *Wolbachia* occurrence as covariates, the quantile-to-quantile plot and the genomic inflation factor (Lambda) was close to 1, indicating that the genomic control of the GWAS was adequate (Fig. S14).

The GWAS identified 14 SNPs significantly associated with the cryptic mtDNA lineage occurrence with *p-values* lower than the Bonferroni adjusted threshold of 4.4 10^-8^ (Fig. 4, S15, Table S11). Out of the 14 SNPs, seven were found close (within 1kb) or within transcripts, and two among them felt within two annotated genes: *SCRASP1 (AGAP005625)* and *CYP6Z1 (AGAP008219).* The gene encoding for the scavenger receptor *SCRASP1* was previously identified in *An. gambiae* as a prominent component involved in immunity response to *Plasmodium* infection, but also other pathogens like bacteria (Danielli, et al. 2000; Christophides, et al. 2002; Stathopoulos, et al. 2014; Smith, et al. 2016). By silencing this gene, Smith, et al. (2016) showed it was an important modulator of *Plasmodium* development in *An. gambiae*. These authors observed that *SCRASP1* was highly enriched after blood-feeding alone and speculated that it may contribute to a metabolic pre-emptive immune response activated by the hormonal changes that accompany blood feeding. Its role as cell surface receptors suggest it may act as immuno-suppressors that when silenced, increase innate immune signaling in mosquito hemocyte populations.

The second genes significantly associated with the cryptic mtDNA haplogroup occurrence was *CYP6Z1*, encoding for a cytochrome P450 capable of metabolizing insecticide like the DDT in *An. gambiae* (Chiu, et al. 2008). *CYP6Z1* is considered more generally as important insecticide resistance gene (Liu 2015; Ibrahim, et al. 2016).

We also identified two additional suggestive mito-nuclear association signals of interest on the X chromosome, with a marginal *p-value* ranging between 5.3 10^-8^ and 1.4 10^-7^ (Fig. 4 and S16). The first one (AGAP000561) is located at *ca.* 9.95Mb and encodes for a Piwi-interacting RNA (piRNA) previously identified as part of “reproductive and development” cluster involved in germline development and maintenance, spermatid development, oogenesis, and embryogenesis of *An. gambiae* (George, et al. 2015). The authors suggested these piRNA plays a significant role in the epigenetic regulation of the reproductive processes in *An. gambiae*. AGAP000561 is an ortholog of the *D. melanogaster* kinesin heavy chain (FBgn0001308), which plays a role in *oskar* mRNA localization to the pole plasm (Brendza, et al. 2000).

The second mito-nuclear marginal association signal of interest on the X chromosome was a clear “skyscraper” located between 15.24Mb and 15.78Mb (Fig. 4). Zooming into this region revealed that the signal contained two skyscrapers with the highest association signals that are nested within a broader region with a distinctive elevation of the *p*-values (see Fig. S16). This distinctive region is not only of special interests for being marginally associated with the cryptic mito-group split, but it was also identified in many populations of *An. gambiae* and *An. coluzzii* with strong signal of recent positive selection and association with metabolic insecticide resistance involving also the mitochondrial oxidative phosphorylation (OXPHOS) respiratory chain (The Ag1000G Consortium 2017; Ingham, Tennessen, et al. 2021; Lucas, et al. 2023) (see also the *Ag1000G Selection Atlas, in prep;* https://malariagen.github.io/agam-selection-atlas/0.1-alpha3/index.html). A total of 32 genes overlap with this focal region (Fig. S16). Among them is the well-known cytochrome p450 encoded by *CYP9K1*, an important metabolic insecticide resistance gene (Main, et al. 2015; Vontas, et al. 2018; Lucas, et al. 2023). Even if that gene is within the elevated *p-value* region, it is located 62kb upstream from the first skyscraper signal. The first highest signal overlapped with AGAP000820 (CPR125 - cuticular protein RR-2 family 125), AGAP00822 and AGAP00823 (CD81 antigen). The second skyscraper was centered close to AGAP000840 (amiloride-sensitive sodium channel) and to AGAP000842 (NADH dehydrogenase (ubiquinone). Other noteworthy genes in that genomic region included include AGAP000849 (NADH dehydrogenase (ubiquinone) 1 beta subcomplex 1), and AGAP0008511 (cytochrome c oxidase subunit 6a, mitochondria). These results thus suggest that the mtDNA lineage haplogroups are associated with mitochondrial genes located in the nuclear genome, as well as genes involved directly or indirectly in insecticide resistances mechanisms (cytochrome p450 and also cuticular regulation genes) and immunity.

Overall, these results support the hypothesis of a tight co-evolutionary history between the two genomic compartments and suggest these mito-nuclear interactions left imprints on the mtDNA genetic variation. The associations between the mtDNA lineages with genes involved in metabolic resistance to pathogens (*Plasmodium* and bacteria) and insecticides support the emerging picture of the key role played by mitochondria, and especially the OXSPHOS pathway in mosquito immunity and insecticide resistance. Previous studies demonstrated that mitochondrial reactive oxygen species (mtROS) produced by the OXSPHOS pathway modulate *An. gambiae* immunity against bacteria and *Plasmodium* (Molina-Cruz, et al. 2008). Ingham, Brown, et al. (2021) were already discussing the disruption of parasite development due to changes in redox state shown experimentally through reducing catalase activity which in turn reduces oocyst density in the midgut (Molina-Cruz, et al. 2008), whilst the initial immune response to parasite invasion consists in a strong mtROS burst (Molina-Cruz, et al. 2008; Castillo, et al. 2017).

Evidence implicating the mitochondrial respiration, OXPHOS pathway, and more generally the mosquito metabolism into metabolic insecticide resistance is increasingly reported in the literature (e.g., Oliver and Brooke 2016; Ingham, Brown, et al. 2021; Ingham, Tennessen, et al. 2021; Lucas, et al. 2023). Ingham, Tennessen, et al. (2021) used a multi-omics study to investigate the causative factors involved in the re-establishment of pyrethroid resistance in a population of *An. coluzzii* colony from Burkina Faso after a sudden loss of the insecticide resistance. Beside the involvement of the 2Rb inversion and of the microbiome composition, the authors detected an increase in the genes expression within the OXPHOS pathway in both resistant populations compared to the susceptible control, which translated phenotypically into an increased respiratory rate and a reduced body size for resistant mosquitoes. This, and previous studies (Oliver and Brooke 2016; Ingham, et al. 2017; Ingham, Brown, et al. 2021), clearly indicated that elevated metabolism was linked directly with pyrethroid insecticide resistance. Additionally, Lucas, et al. (2023) investigated novel loci associated with pyrethroid and organophosphate resistance in *An. gambiae* and *An. coluzzii* using a GWAS, which also implicated the involvement of a wide range of cytochrome p450, mitochondrial, and immunity genes (including also the same genomic region on the X chromosome as the one we detected here). Both Ingham, Tennessen, et al. (2021) and Lucas, et al. (2023) further pointed out possible cross-resistance mechanisms in metabolic insecticide resistance at large, in which the mosquito metabolism, mitochondrial respiration, the OXPHOS pathway, and mtROS production all seem to play a central role.

## Conclusions

In this study we showed that the determinants of mitochondrial genetic variation are multifarious and complex. In agreement with previous studies (Fontaine, et al. 2015; Thawornwattana, et al. 2018; Müller, et al. 2021), the mtDNA phylogeny clearly illustrated the previously reported highly reticulated evolutionary history of the AGC. On one side, the three most widely distributed species – *An. gambiae*, *An. coluzzii*, and *An. arabiensis* – form a rather homogeneous mtDNA gene pool clearly illustrating the extensive level of introgression that occurred between them over the evolutionary timescale of the AGC. On the other side, other species of the AGC cluster in a well diverged monophyletic clade, where each species forms a clearly distinct monophyletic group. One haplotype in the mtDNA gene pool of *An. gambiae* / *An. coluzzii* clustered close to *An. quadriannulatus,* in a position of the species tree where *An. arabiensis* was placed according to the species informative loci on the X chromosome and the autosomes (Fontaine, et al. 2015). This suggest that *An. arabiensis* mito-lineage might still be segregating in this joined mtDNA gene pool of the three most widespread species in the AGC. Mitochondrial introgression was also detected in other species, notably between *An. bwambae* and most likely *An. gambiae*. The mitochondrial phylogenetic clustering of all the salt-water tolerant members of the AGC (*An. merus, An. melas*, and *An. bwambae)* into a strongly supported monophyletic group also departed from the admitted species branching order (Fontaine, et al. 2015; Thawornwattana, et al. 2018; Barrón, et al. 2019). This suggests that historical mtDNA capture or selection from ancestral standing genetic variation may have occurred, possibly involving selective processes related to specialization to a very distinct salty larval habitat compared to the other fresh-water tolerant species of the AGC, and to the majority of the *Anophelinae* species (Bradley 1994; Bradley 2008). A proper population genetic study investigating this specialization from an evolutionary perspective still remains to be done.

Population structure, demography, and dispersal were found to be key drivers shaping the mtDNA variation across the African populations of *An. gambiae* and *An. coluzzii*. The patterns identified mostly followed those previously reported at the nuclear genomes (The Ag1000G Consortium 2017, 2020). Despite the extensive level of gene flow between *An. gambiae* and *An. coluzzii*, significant variance partitioning between species was still detectable. Even more striking was a clearly distinct mito-lineage composed of 244 samples from *An. gambiae* and *An. coluzzii*. This lineage displayed a level of divergence comparable to, yet slightly lower than, the mtDNA divergence observed between the species of the AGC. Its distribution closely matched the distribution of *An. coluzzii* with a West-to-East and North-to-South decreasing frequency gradient. This cryptic lineage appears to be a phylogeographic legacy of the species isolation followed by a secondary contact between *An. gambiae* and *An. coluzzii* with incomplete homogenization. Its frequency was clearly associated with species divergence (being enriched in *An. coluzzii* compared to *An. gambiae*), mitochondrial level of diversity, and with population structure, but it was not linked with the rare *Wolbachia* occurrence detected from the short-read data. Once accounting for these variables in a GWAS-like study, we found significant associations between the cryptic lineage occurrence and SNPs of the nuclear genome mostly from genes involved in metabolic resistance to pathogens and insecticides. These results suggest that the phylogeographic split of mitochondrial lineages and its incomplete re-homogenization after the secondary contact involved selective processes and a certain mito-nuclear coevolution process between the two genome compartments. These associations support the picture emerging in the recent literature underlining the key role played by the respiratory metabolism, the OXPHOS pathway, and the generation of reactive oxygens in the metabolic resistance to pathogens and to insecticides.

Cross-resistance mechanisms are increasingly recognized as a major threat to vector control strategy allowing mosquitoes to adapt to insecticides (Ingham, Tennessen, et al. 2021; Lucas, et al. 2023). Our results call for additional studies characterizing further the extent of mito-nuclear associations, the role of mitochondria in adaptive processes to pathogens and insecticides, and a better understanding of the tight coordination and co-evolution between the mitochondrial and nuclear genome. By integrating both mtDNA and nuclear data, this study underlines that the mtDNA locus, once considered as a nearly neutral locus and thus informative on the phylogenetic history of species, has in fact a much more complex evolution in *Anopheles* mosquitoes where all the evolutionary forces (drift, migration, mutation and multiple type of selection) interact. Such integration of nuclear and mito-genomic study are still rare, but necessary to further our understanding of insect genomic evolution (Cameron 2014).

## Materials and methods

### Sampling and whole genome short read data

We retrieved whole genome short read (WG-SR) data (100 bp paired-end Illumina sequencing) from 74 mosquito specimens for six species of the AGC from Fontaine, et al. (2015), including *An. gambiae s.s, An. coluzzii, An. arabiensis, An. quadriannulatus, An. melas, and An. merus*. As mtDNA genomes of these samples were previously assembled, we compared them with the ones produced using the new pipeline developped in the present study. We extracted reads that did not map to the nuclear reference genome and used them to assemble mitogenome. We included also WG-SR data from three specimens of a seventh species – *An. bwambae –* that were generated as part of the Anopheles 16 Genomes Project (Fontaine, et al. 2015; Neafsey, et al. 2015). See the complete sampling details in Figure S1 and Table S1. We also retrieved WG-SR data from The Ag1000G Consortium (2020) phase 2-AR1 release consisting of 1142 wild-caught mosquito specimens including *An. gambiae s.s.* (n=720), *An. coluzzii* (n=283) and hybrid (n=139) from 16 geographical sites (Figure 1, Table S2).

Previously generated *An. gambiae* reference mitochondial genome (GenBank ID: L20934.1) (Beard, et al. 1993) was used to guide the assembly of the *AutoMitoG* pipeline. The mitochondrial sequences of *An. christyi* and *An. epiroticus* from Fontaine, et al. (2015) were also included as outgroups sequences for phylogeographic and phlogenetic analyses.

### Mitochondrial genomes assembly and alignment

Information about the software versions is provided in Table S3. From the WG-SR files mapped to nuclear reference genomes (bam files) obtained from Fontaine, et al. (2015) and The Ag1000G Consortium (2020), we extracted reads that did not mapped to the nuclear reference genome and converted them to paired-reads fastq files using *Samtools* (Li, et al. 2009) and *Picard* Tools (http://broadinstitute.github.io/picard/). We wrote the *AutoMitoG* [*Automatic Mitochondrial Genome assembly*] pipeline to streamline the mitochondrial genome assembly process (available at https://github.com/jorgeamaya/automatic_genome_assembly, Fig. S2). As a general overview, the pipeline starts by randomly sampling paired reads from each file at a 5% rate. Then, the pipeline proceeds to assemble the mitogenome using a modified version of MITObim (Hahn, et al. 2013) (see details below) and evaluate the quality of the assembly. This is done by counting the number of ambiguities outside the *D-loop* control region; a region prone to sequencing and assembly errors due to the AT-rich homopolymer sequences. If ambiguities remain in the mtDNA assembly, the previous steps are repeated iteratively, increasing the sampling rate by 5% until the assembled mitogenomes shows no ambiguities or until 100% of the reads are used. If ambiguities persist after reaching a sampling rate of 100%, the assembly with the least number of ambiguities is selected by default. Finally, assembled mitogenome sequences together with previously assembled reference genome (Beard, et al. 1993), and outgroup mtDNA sequences were aligned to each other with MUSCLE (Edgar 2004).

The *AutoMitoG* pipeline relies on a modified version of MITObim (Hahn, et al. 2013) to assemble the mtDNA genomes. Subsampling of the paired-reads is performed to achieve two purposes: (1) minimize the number of ambiguous base calls – these result from conflicting pairing of reads from mitochondrial origin with reads possibly originating from nuclear mitochondrial DNA copies (NUMTs), reads with sequencing errors, and reads that originated from possible contamination. Since the number of reads from mitochondrial origin is orders of magnitude larger in the WG-SR data than the number of reads from other sources, subsampling safely reduces offending reads; (2) to normalize the dataset coverage, which speeds up MITObim calculations, as the proportion of mitochondrial reads can differ between samples and studies. Indeed, MITObim performs best with sequencing depth between 100 and 120x for Illumina reads (Hahn, et al. 2013).

MITObim performs a two-steps assembly process (see Fig. 2 in Hahn, et al. (2013)). First, it maps reads to a reference genome, here the mtDNA genome of *An. gambiae* from Beard, et al. (1993), to generate a “backbone”; second, it extends this “backbone” with overlapping reads in an iterative *de novo* assembly procedure. Thanks to its hybrid assembly strategy, MITObim perform well even if the samples and the reference genome are phylogenetically distant (Hahn, et al. 2013). We forced majority consensus for non-fully resolved calls during the backbone assembly and during the backbone iterative extension, for which we customized MITObim’s code. The original version of MITObim does not force majority consensus and was not used in this study. However, it is included as an option in the pipeline for the benefit of users who may prefer less stringent assembly criteria. See Fig. S2 for further information on the pipeline usage and the corresponding documentation in the GitHub page.

We compare the newly assembled mitogenomes with those previously generated in Fontaine, et al. (2015) (n=74). For that purpose, we first aligned mitogenome sequences from the two studies, cropped out the control region (sequence length=14,844bp) following Fontaine, et al. (2015) as it is prone to sequencing and assembly errors. Then, for each pair of mtDNA assemblies (new vs previous) coming from each of the 74 samples in Fontaine, et al. (2015), we counted the number of pairwise differences. We also visually compared assemblies generated with the two pipelines by building a distance-based neighbor-joining tree (HKY genetic distance model) (Fig. S4). These steps were conducted in Geneious Prime® (2023.0.1, Build 2022-11-28 12:49).

### Mitogenome genetic diversity and phylogenetic relationships

As an initial assessment of the mtDNA alignment characteristics, we calculated various estimators of genetic diversity per species and per location including: the number INDEL sites, segregating sites (S), average number of differences between pairs of sequences (K), number of haplotypes (H), haplotype diversity (HD), nucleotide diversity (π), Theta-Waterson (Θ_W_), the Tajima’s D, and the Achaz’s Y (Achaz 2008). These statistics were computed using the C-library *libdiversity* developed by G. Achaz (https://bioinfo.mnhn.fr/abi/people/achaz/cgi-bin/neutralitytst.c).

We estimated the phylogenetic relationships among mtDNA haplotypes using the same methodology as in Fontaine et. al. (2015). We constructed mtDNA maximum likelihood (ML) phylogenies using RAxML (parameters -m GTRGAMMA -# 1000 -T 16 -f a -x 12345 -p 12345). Bootstrap nodes’ supports were calculated using the fast-bootstrap method of RAxML (T 16 -# 1000 -f b -m GTRGAMMA). The mitogenome sequences from *An. christy* and *An. epiroticus* were used as outgroups to root the trees. Multiple ML phylogenetic trees were built: one only considering the 74 sequences from the *An. gambiae* species complex for comparative purpose with previously published ML tree in Fontaine, et al. (2015), including also the 3 *An. bwambae* samples; and another tree considering all the mitogenome sequences including the 77 mitogenome sequences combined with the 1142 sequences of *An. gambiae* and *An. coluzzii* samples from The Ag1000G Consortium (2020). To ease visualization, clades were collapsed when possible if they contained multiple closely related samples.

In order to provide an alternative visualization of the phylogenetic relationships given the large size of the total alignment, we also visualized mtDNA genetic variation among the 1142 sequences of *An. gambiae* and *An. coluzzii* samples from The Ag1000G Consortium (2020) into a reduced multidimensional space using a non-metric multidimensional scaling (nMDS). For that purposed, we calculated a *p*-distance matrix among sequences using MEGA v.7 (Kumar, et al. 2016) and performed the nMDS using the *ecodist* R-package (Goslee and Urban 2007). The nMDS results were further processed using *scikit*-learn v.0.22.1 (Pedregosa, et al. 2011) to identify major clusters in the data set, using a hierarchical clustering algorithm.

### MtDNA genetic structure in natural populations of *An. gambiae* and *An. coluzzii*

We first assessed how the mtDNA variation of 1142 sequences of *An. gambiae* and *An. coluzzii* samples from The Ag1000G Consortium (2020) partitioned among different levels of structuration using an analysis of molecular variance (AMOVA) (Excoffier, et al. 1992). We considered 3 hierarchical levels of stratification: between species, among populations within species, and within populations. The AMOVA was conducted with the TN93+gamma model of sequence evolution using the *poppr R-package* (Kamvar, et al. 2014) and the AMOVA function derived from the APE v5.6-4 R-package (Paradis, et al. 2004; Paradis and Schliep 2019). Significance test was conducted using 1000 permutations. The analysis was conducted considering only populations from the Ag1000G that were taxonomically unambiguous (n=938, see Fig. 1), thus removing the hybrid taxonomic uncertain populations from The Gambiae (GMS) and Guinea-Bissau (GWA), as well as the taxonomically uncertain population from Kenya (KEA).

Then, we quantified the level of genetic differentiation among populations by calculating the pairwise *F_ST_* differences using Arlequin v3.5 (Excoffier and Lischer 2010). We compared *F_ST_* values obtained for the mitochondrial DNA (mtDNA) with those previously reported for the nuclear genome (nDNA) (The Ag1000G Consortium 2020).

We characterized further the mtDNA variation, comparing genetic diversity estimators for each species at each locality. Since sample sizes vary among locations and can influence diversity estimators, we performed a rarefaction procedure to account for differences in sample sizes (Hurlbert 1971; Kalinowski 2004, 2005; Szpiech, et al. 2008; Colwell, et al. 2012). To do so, sequences from each location were randomly sampled incrementally, starting with three sequences up to a maximum of 50 sequences or until there were no more sequences available for the specific location. This random sub-sampling with replacement of the sequences was repeated 5000 times for each sample size increment (from 3 to 50) to estimate the mean and standard error of the statistic of interest. This rarefaction analysis was applied for estimating the standardized number of segregating site (*S*), the number of haplotypes (*H*), the nucleotide diversity (*π*), the Tajima’s *D*, and the Achaz’s *Y* using python scripts and the *c-library libDiversity*. Results for each statistic were summarized as rarefaction curves.

Isolation-by-distance (IBD) was computed following Rousset (1997). We derived the unbounded level of genetic differentiation *F_ST_*/(1-*F_ST_*) between pairs of populations and correlated the genetic distance with the geographic distance, expressed as the great circle distance (in log_10_ unit) globally across species, and also for each species separately. The strength and significance of the IBD was tested using a Mantel test implemented in *ade4* R-package (Dray and Dufour 2007) with 1000 permutations of the geographic distance matrix. Since we were interested only in testing IBD within well-defined species, we removed the hybrid and taxonomically ambiguous populations (The Gambia – GM, Guinea Bissau – GW, and Kenya – KEA) from this analysis. Likewise, we ran the analysis with and without the island *An. gambiae* population of Mayotte (FRS), as this population departs from the species’ continuum (The Ag1000G Consortium 2020).

### Detection of *Wolbachia* infection in natural populations of the Ag1000G

MtDNA variation can be strongly impacted by cytoplasmic conflict with the endosymbiont Wolbachia (Galtier, et al. 2009; Dong, et al. 2021), and this latter has been reported in the AGC (Baldini, et al. 2014; Shaw, et al. 2016; Gomes, et al. 2017; Gomes and Barillas-Mury 2018; Jeffries, et al. 2018; Ayala, et al. 2019; Chrostek, et al. 2019; Straub, et al. 2020; Jeffries, et al. 2021). Therefore, we used the unmapped WG-SR data to diagnose the infection status of each mosquito specimens of *An. gambiae* and *An. coluzzii* from the Ag1000G phase-II (The Ag1000G Consortium 2020). To that end, we screened the unmapped Ag1000G WG-SR data to detect *Wolbachia* specific sequences using MagicBlatst v.1.1.5 (NCBI) (Boratyn, et al. 2019) following the procedure described in Pascar and Chandler (2018). WG-SR reads that did not map to the nuclear reference genome were compared to selected reference *wsp*, *ftsZ*, and *groE* operon sequences isolated from *Wolbachia* samples that are representative of supergroups A to D. For our analysis, we used the *Wolbachia* sequence database of Pascar and Chandler (2018), which includes 61 sequences of *Wolbachia* type A to D. We added four new sequences assembled by the authors to their database. These are *Wolbachia* sequences of type B also found in *An. gambiae* specimens (Pascar & Chandler 2018). Using the same *(strict)* detection criterion as in Pascar and Chandler (2018), a minimum of three reads with at least 98bp length and 95% identity had to match with the same *Wolbachia* sequences for the specimen to be considered as infected. We also applied a more “*lenient”* criterion: a minimum of three reads with at least 90bp length and 90% identity had to match with the same *Wolbachia* sequences for the specimen to be considered infected.

### Mitogenome lineages associations with genomic features and with SNPs of the nuclear genome

We explored the associations between the two main mitochondrial phylogenetic lineages and the genetic variation on the nuclear genome using the genome-wide SNP data from The Ag1000G Consortium (2020). We also considered the association of the mtDNA lineages with other covariates including population structure, *Wolbachia* infection status, and major chromosomal inversions. For that purpose, we used a genome wide-like association study (GWAS) (Ansari, et al. 2017; Fellay and Pedergnana 2019), considering the two main mtDNA lineages discovered in the phylogenetic analyses and how they are associated with each SNP in the nuclear genome. This design aimed to assessing the extent of functional associations between mtDNA lineages and the nuclear genome, highlighting potential mito-nuclear co-evolution history, considering covariates, such as populations structure and *Wolbachia* infection status.

Following standard practices in GWAS (Uffelmann, et al. 2021), we first ensured that the samples included in the Ag1000G were not too closely related. Therefore, we estimated the within population kinship coefficients using KING 2.2.4 (Manichaikul, et al. 2010). The KING-robust approach relies on relationships inference using high density SNP data to model genetic distance between pairs of individuals as a function of their allele frequencies and kinship coefficient (Manichaikul, et al. 2010). This contrast with other methods such as the one implemented in PLINK (Purcell, et al. 2007) which estimates relatedness using estimator of pairwise identity-by-descent. However, this method is very sensitive to population demography in contrast to KING-robust approach.

Following The Ag1000G Consortium (2017, 2020), we only used the free-recombining biallelic SNPs (n=1,139,052) from section 15M to 41M of chromosome 3L to estimate pairwise kinship coefficients. This genomic portion avoids non-recombining centromeric regions and major polymorphic chromosomal inversions on chromosome 2, and the sex chromosome. No down sampling, nor linkage disequilibrium (LD) pruning, nor any other preprocessing was undertaken on the data, following KING’s authors recommendation (Manichaikul, et al. 2010). We iteratively removed individuals with the largest number of relationships above 2^nd^ degree relative as estimated by KING. At any step, when two individuals were found to have the same number of relationships, we removed the first individual according to its identifier’s alpha numeric order.

In order to include population structure as a covariate in the GWAS analysis, we conducted a principal component analyses (PCA) following the same procedure as described in The Ag1000G Consortium (2017, 2020). We selected randomly 100,000 biallelic SNPs from the free-recombining part of the genome of *An. gambiae* and *An. coluzzii* on chromosome 3L (from position 15Mb to 41Mb). To remain consistent with The Ag1000G Consortium (2017, 2020), we followed the same filtering procedure. We performed a LD-pruning using the function *locate_unlinked* from Python’s module *scikit-allel* version 1.2.1 (Miles and Harding 2016) to ensure independence among SNPs. Specifically, we scanned the genome in windows of 500bp slid by steps 200bp and excluded SNPs with an *r^2^* ≥ 0.1. This process was repeated 5 times to ensure most SNPs in LD were removed. A PCA was then performed as described in The Ag1000G Consortium (2017, 2020), using *scikit-allel* version 1.2.1 (Miles and Harding 2016). Results were plotted highlighting specimens according to their locality of origin. PC scores were stored and used as covariates in the GWAS analysis.

Prior to the actual GWAS analysis, we assessed the extent of association between the two identified major mtDNA phylogenetic lineages, the specimens’ sex, *Wolbachia* infection status, and population structure as estimated by the top six PC axes from the PCA. We also considered the inversion karyotypes as reported by The Ag1000G Consortium (2020) and (Love, et al. 2019). Given the diverse nature of covariables, we calculated a proxy of the Pearson’s correlation coefficient between variables capable of handling numerical and categorical variable types using the *x2y*-metric (Ramakrishnan 2021; Lares 2023). The *x2y*-metric performs a linear regression on continuous response variables and a classification procedure on categorical response variables. Then, it uses the calculated model to predict the data based on the independent variable and, finally, estimates a percentage of error in the predictions. As this method does not provide any significance test with a *p*-value, the 95% confidence interval was calculated using 1000 permutations.

Finally, we conducted the formal GWAS-like analysis to evaluate the associations between the two main mtDNA phylogenetic lineages and the nuclear SNP genotype variation, considering the following covariates: the population structure using PC-scores, the *Wolbachia* infection status, and sex. We performed the GWAS using the program *SNPTEST* v2.5.4-beta3 (Marchini *et al*. 2007). The main mtDNA lineage were used as “phenotype values” defined as the phylogenetic mtDNA clusters from the hierarchical clustering of the NMDS analysis. After normalization, we used the PC scores of the PCA obtained from S*cikit-allele* as continuous covariates, *Wolbachia* infection status and sex as binary covariates. Only unrelated samples (n=1053) and SNPs with a MAF ≥ 0.01 (n=7,858,575, Table 10) were considered in this analysis. The threshold to assess the significance of the GWAS was defined following a Bonferroni corrected *p*-value accounting for the number of independent genomic blocks in the genome (0.05/1,139,052 = 4.39e-8). The number of independent genomic blocks in our data set was approximated by the number of independent SNPs as determined with Plink v 1.90 (Table S10). The results of the GWAS were plotted as Manhattan and QQ-plots in R.

We produced a list of genes IDs that contained one or more significant SNPs from the GWAS within the CDS or within 1kb upstream or downstream from the CDS according to the general feature format file VectorBase-57_AgambiaePEST.gff from *VectorBase* release 57, 2022-APR-21 (Giraldo-Calderon, et al. 2015). The list of genes was then used to extract information associated to such genes from *VectorBase*.

## Acknowledgment

We would like to acknowledge the *Anopheles gambiae* 1000 genomes consortium (https://www.malariagen.net/projects/ag1000g#people), and especially Nick Harding, Mara K.N. Lawniczak, Martine Donnelly, Dominic P. Kwiatkowski (deceased), Carlo Costantini, Nora J. Besansky, and Frédéric Labbé for providing resources, supports, and help during the development of this study. We also thank the Center for Information Technology of the University of Groningen for their support and for providing access to the *Peregrine* high-performance computing cluster. We also wish to acknowledge the ISO 9001 certified IRD *i-Trop* HPC (member of the South Green Platform) at IRD Montpellier for providing HPC resources that have contributed to the research results reported within this paper (https://bioinfo.ird.fr; www.southgreen.fr). This study was supported by the University of Groningen (The Netherlands) through a phD fellowship of the Adaptive Life program awarded to JEAR and through a starting grant awarded to MCF. CC was supported by a GAIA PhD fellowship from the University of Montpellier (France).

## Author contribution

MCF designed research; JEAR and MCF performed research; AM, CC, YBC, VP contributed new data/reagents/analytic tools; AM and CC curated the original data of the Ag1000G phase-2; JEAR, MCF, YBC, CC analyzed data; MCF and BW provided supervision and funding; MCF wrote the paper; all the authors provided inputs and feedbacks.

## Data availability

Short-read data used in this study to assemble mitogenomes come from two consortium projects: the MalariaGEN *Anopheles gambiae* 1000 genomes projects phase-2 AR1 data release (https://www.malariagen.net/data/ag1000g-phase-2-ar1; see also The Ag1000G Consortium (2020)); and the Anopheles 16 genomes project (Fontaine, et al. 2015; Neafsey, et al. 2015) (National Center for Biotechnology Information, NIH, BioProject IDs: PRJNA67511 and PRJNA254046). Mitochondrial genome sequences have been deposited on NCBI and the mitogenome alignment on the IRD DataSud repository: https://doi.org/10.23708/XXXXX. The *AutoMitoG* pipeline, codes and scripts used in this study are available via github (https://github.com/jorgeamaya/automatic_genome_assembly and https://github.com/jorgeamaya/malaria_mitogenome).

