## Supplementary material for "Mitochondrial variation in *Anopheles gambiae* and *An. coluzzii*: phylogeographic legacy of species isolation and mito-nuclear associations with metabolic resistance to pathogens and insecticides": SI Files

**Running title**: Mitogenome evolution in the *Anopheles gambiae* complex


Jorge E. Amaya Romero[1,2]†, Clothilde Chenal[2,3], Yacine Ben Chehida[1,4], Alistair Miles[5], Chris S. Clarkson[5], Vincent Pedergnana[2], Bregje Wertheim[1], Michael C. Fontaine[1,2]*†

Affiliations

1. Groningen Institute for Evolutionary Life Sciences (GELIFES), University of Groningen, Nijenborgh 7, 9747 AG Groningen, Netherlands.
2. MIVEGEC, Univ. Montpellier, CNRS, IRD, Montpellier, France.
3. Institut des Science de l'Evolution de Montpellier, Univ Montpellier, CNRS, Montpellier, France
4. Ecology and Evolutionary Biology, School of Biosciences, University of Sheffield, Sheffield, S10 2TN, UK
5. Wellcome Sanger Institute, Hixton, Cambridge CB10 1SA, UK

† contributed equally to the study.

*** Correspondance :** Michael C. Fontaine. MIVEGEC (U. Montpellier, CNRS, IRD), Centre IRD Occitanie, 911 Avenue Agropolis, BP 64501, 34394 Montpellier Cedex 5, France

ORCID

- YBC : https://orcid.org/0000-0001-7269-9082
- AM : https://orcid.org/0000-0001-9018-4680
- VP : https://orcid.org/0000-0002-7852-5339
- BW : https://orcid.org/0000-0001-8555-1925
- MCF : https://orcid.org/0000-0003-1156-4154

### Supplementary figures

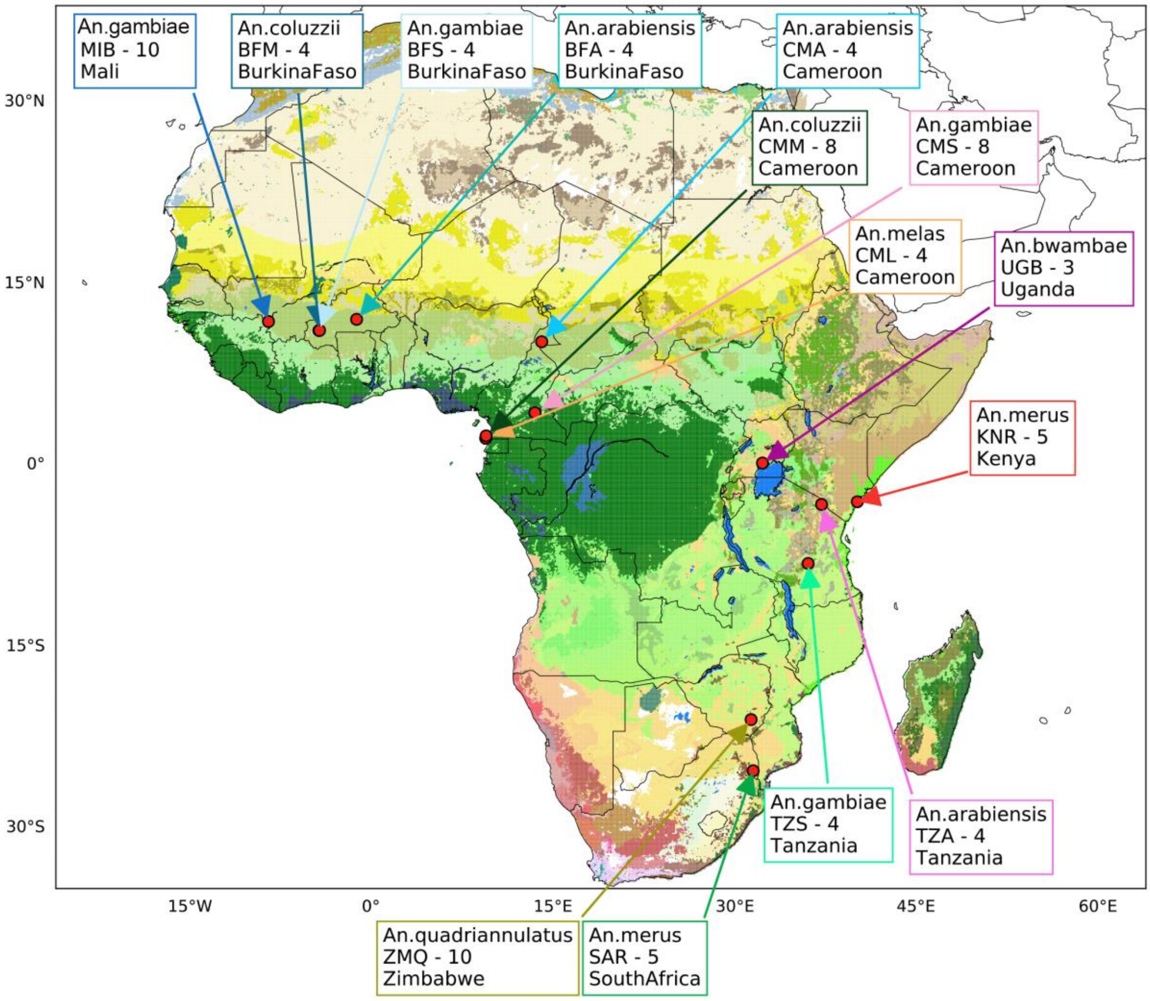


Figure S1. Sampling distribution of seven species of the *An. gambiae* species complex from Fontaine, et al. (2015).

Approximate sampling locations of the 6 species of the *An. gambiae* species complex from Fontaine, et al. (2015), augmented with *3 An. bwambae* samples produced as part of the 16 Anopheles species genome project (Fontaine, et al. 2015; Neafsey, et al. 2015). Red dots show the approximate sampling locations per geographic site. The sampling code (3 letters) as well as the sample size per location is provided, and the country of origin (see **Table S1** for further details on the sampling).

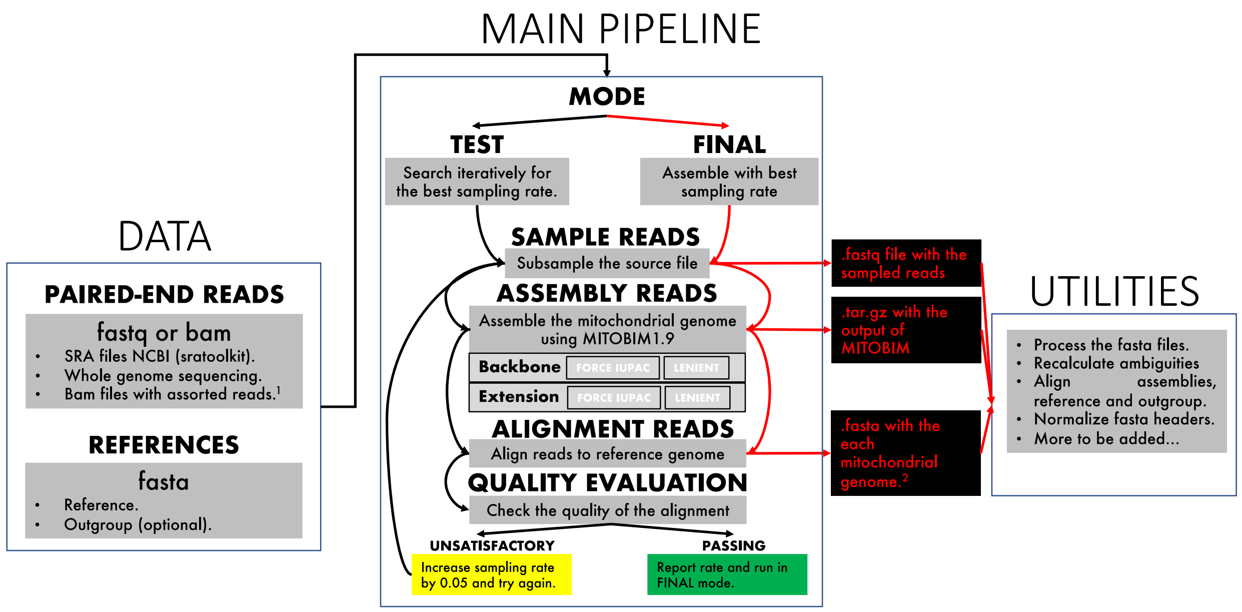


Figure S2. AutoMitoG [*Automated mitochondrial genome assembly*] pipeline workflow.

Unmapped paired-end short read data (fastq or BAM files) are provided together with a mitochondrial reference sequence which can be the entire the mitogenome (if available) or a fragment of it. The pipeline starts by randomly sampling paired reads from each file at a 5% rate. This sub-sampling step is performed to achieve two goals: (1) minimize the number of ambiguous base calls which can result from conflicting pairing of reads from mitochondrial origin with reads possibly originating from nuclear mitochondrial DNA copies (NUMTs), reads with sequencing errors, and reads that originated from possible contamination. Since the number of reads from mtDNA origin is orders of magnitude larger than the number of reads from other sources, sub-sampling safely reduces offending reads; (2) to normalize the dataset coverage, which speeds up MITObim calculations, as the proportion of mtDNA reads can differ between samples and studies. Indeed, MITObim performs best with sequencing depth between 100 and 120x for Illumina reads (Hahn, et al. 2013). Then, the pipeline proceeds to assemble the mitogenome using a modified version of MITObim and evaluate the quality of the assembly. If ambiguities persist in the mtDNA assembly, the previous steps are repeated iteratively, increasing the sampling rate by 5% until the assembled mitogenomes shows no ambiguities or until 100% of the reads are used. If ambiguities persist after reaching a sampling rate of 100%, the assembly with the least number of ambiguities is selected by default. MITObim performs a two-steps assembly process (see Fig. 2 in Hahn, et al. (2013)). First, it maps reads to a reference genome (here the mtDNA genome of *An. gambiae* from Beard, et al. (1993)) to generate a "backbone"; second, it extends this "backbone" with overlapping reads in an iterative *de novo* assembly procedure. Thanks to its hybrid assembly strategy, MITObim perform well even if the samples and the reference genome are phylogenetically distant (Hahn, et al. 2013). We forced majority consensus for non-fully resolved calls during the backbone assembly and during the backbone iterative extension, for which we customized MITObim's code. The original version of MITObim does not force majority consensus and was not used in this study. However, it is included in the pipeline for the benefit of users who may prefer less stringent assembly criteria. See the main text and the GitHub page for further information on the pipeline usage (https://github.com/jorgeamaya/malaria_mitogenome).

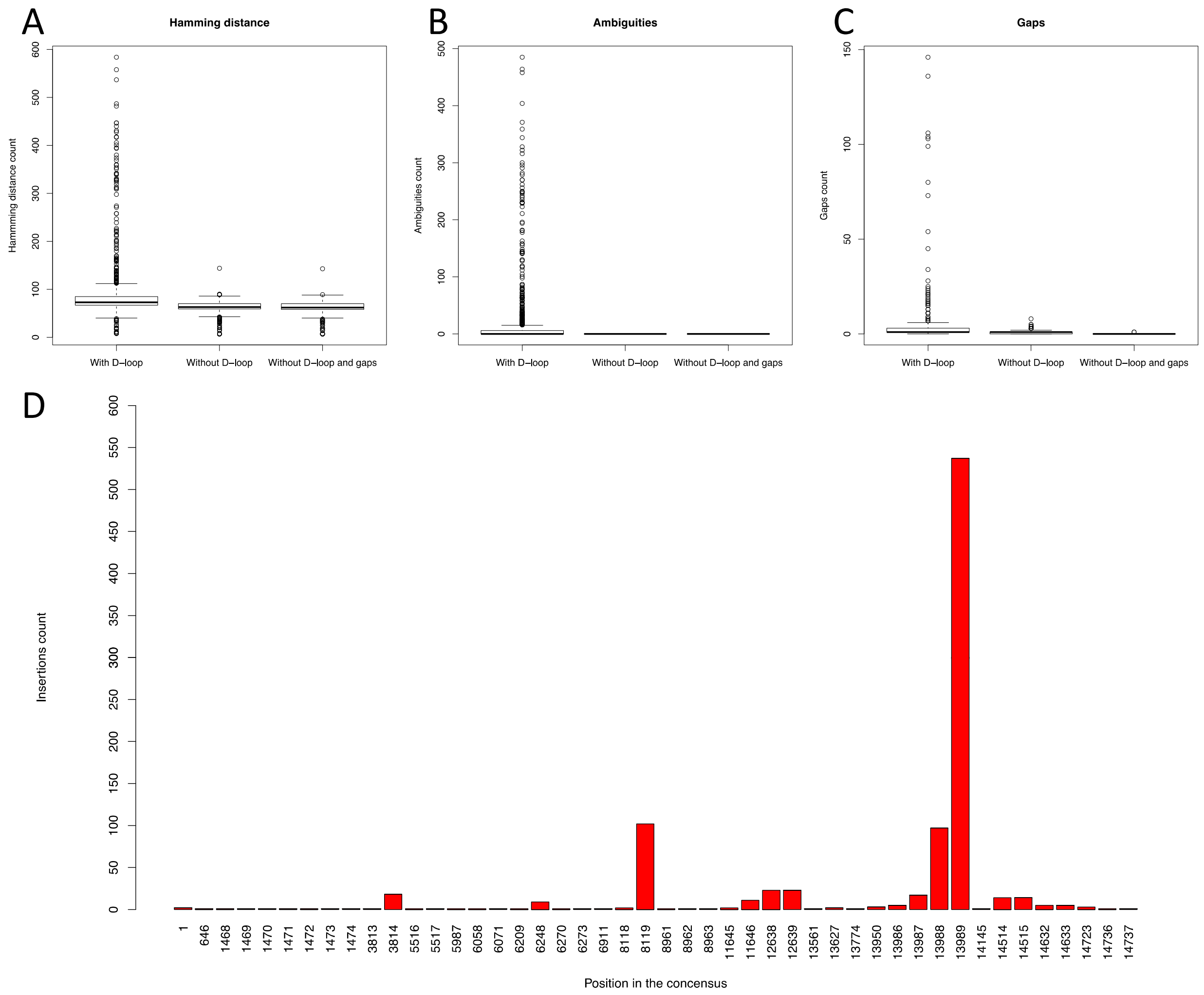


**Figure S3. Quality assessment of the re-assembled mitogenome sequence alignment from Fontaine et al. (2015).**

Comparison of alignments with D-loop (including gaps and the D-loop), without D-loop (including gaps but no D-loop), and without D-loop and gaps (following Fontaine et. al. 2015). **A.** Hamming distance to the reference sequence of Beard et al. (1993) per window of 100bp. **B.** Ambiguities count. **C.** Gaps count. **D**. Position of the gaps and number of sequences in which the gap was filled with a nucleotide.

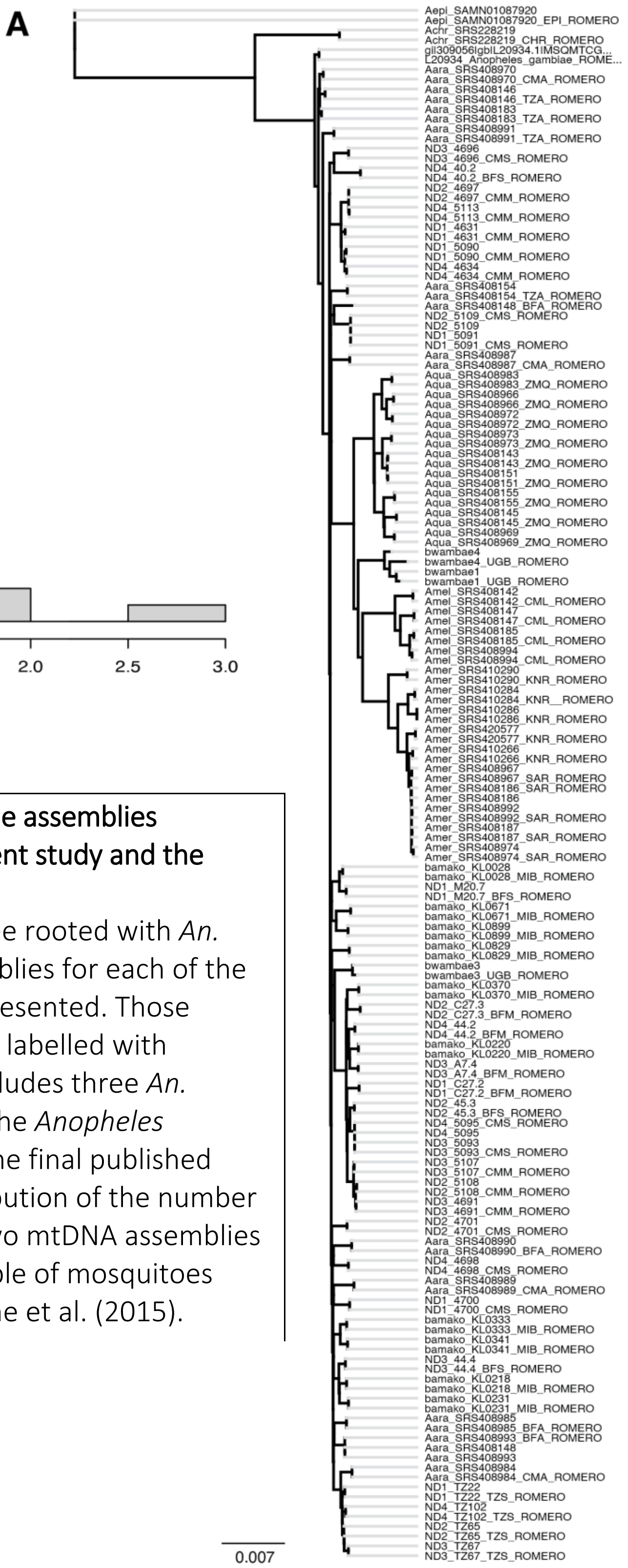


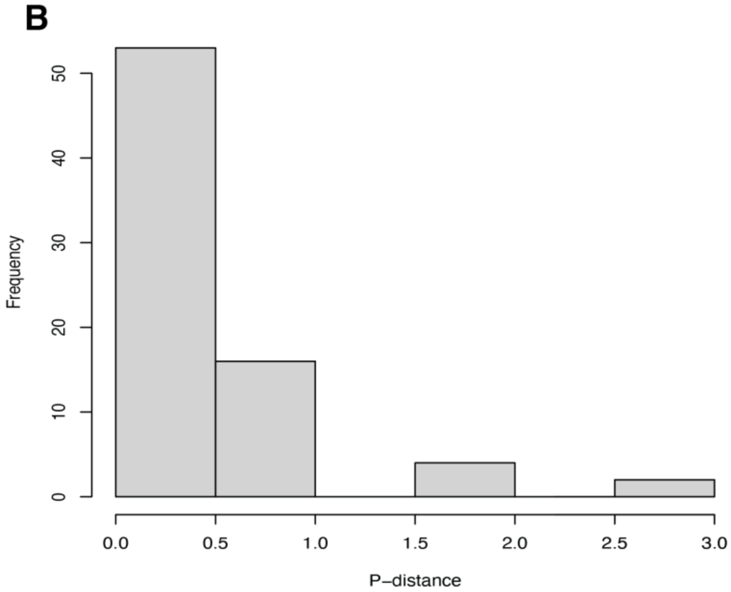


**Figure S4. Comparisons of the mitogenome assemblies generated by the pipelines from the present study and the former one of Fontaine et al. (2015).**
**(A)** Neighbor-Joining *p*-distance based tree rooted with *An. epiroticus* and *An. christyi*. Pairs of assemblies for each of the samples in Fontaine et al. (2015) are represented. Those corresponding to the new assemblies are labelled with "_ROMERO" at the end. The tree also includes three *An. bwambae* samples generated as part of the *Anopheles gambiae* cluster consortium, but not in the final published paper. **(B)** Histogram reporting the distribution of the number of nucleotide differences between the two mtDNA assemblies generated for each sample for each sample of mosquitoes from the *An. gambiae* complex in Fontaine et al. (2015).

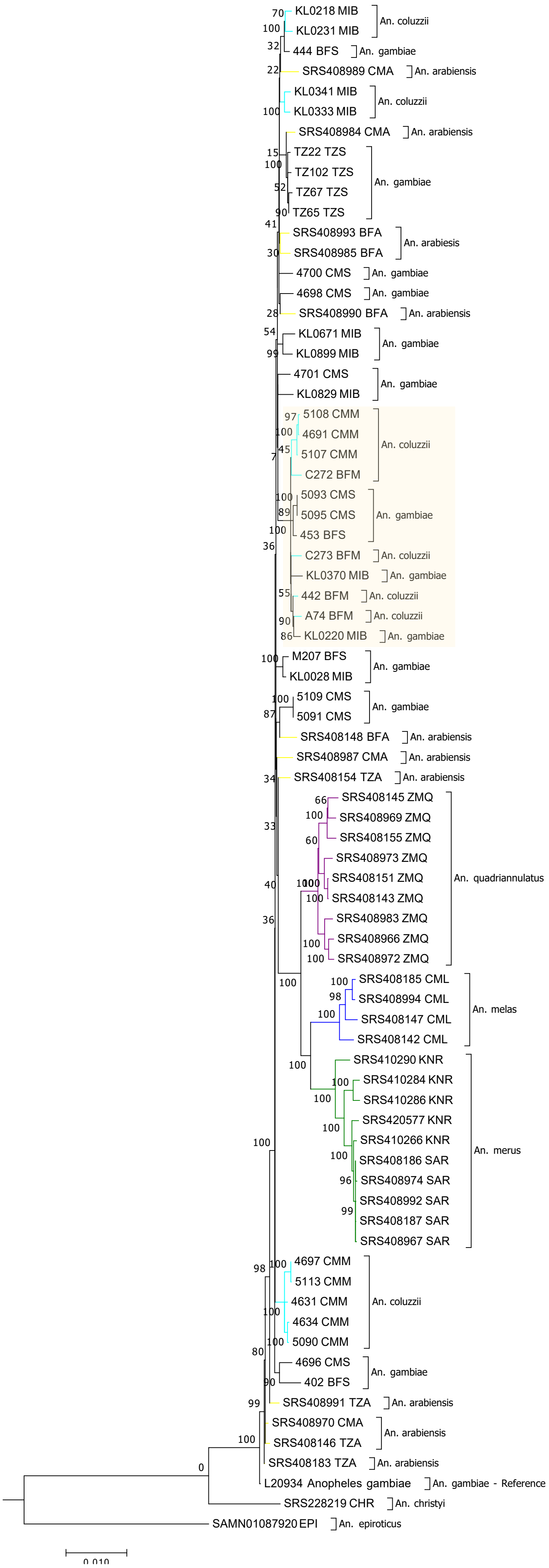


**Figure S5.** **RAXML maximum likelihood phylogeny for the mitochondrial genomes newly re-assembled in this study for 74 samples of the *An. gambiae* complex (or AGC) from Fontaine, et al. (2015).** Bootstrap support values (%) are indicated at the nodes. The tree is rooted mtDNA sequences from two outgroup species: *An. christyi* and *An. epiroticus*.

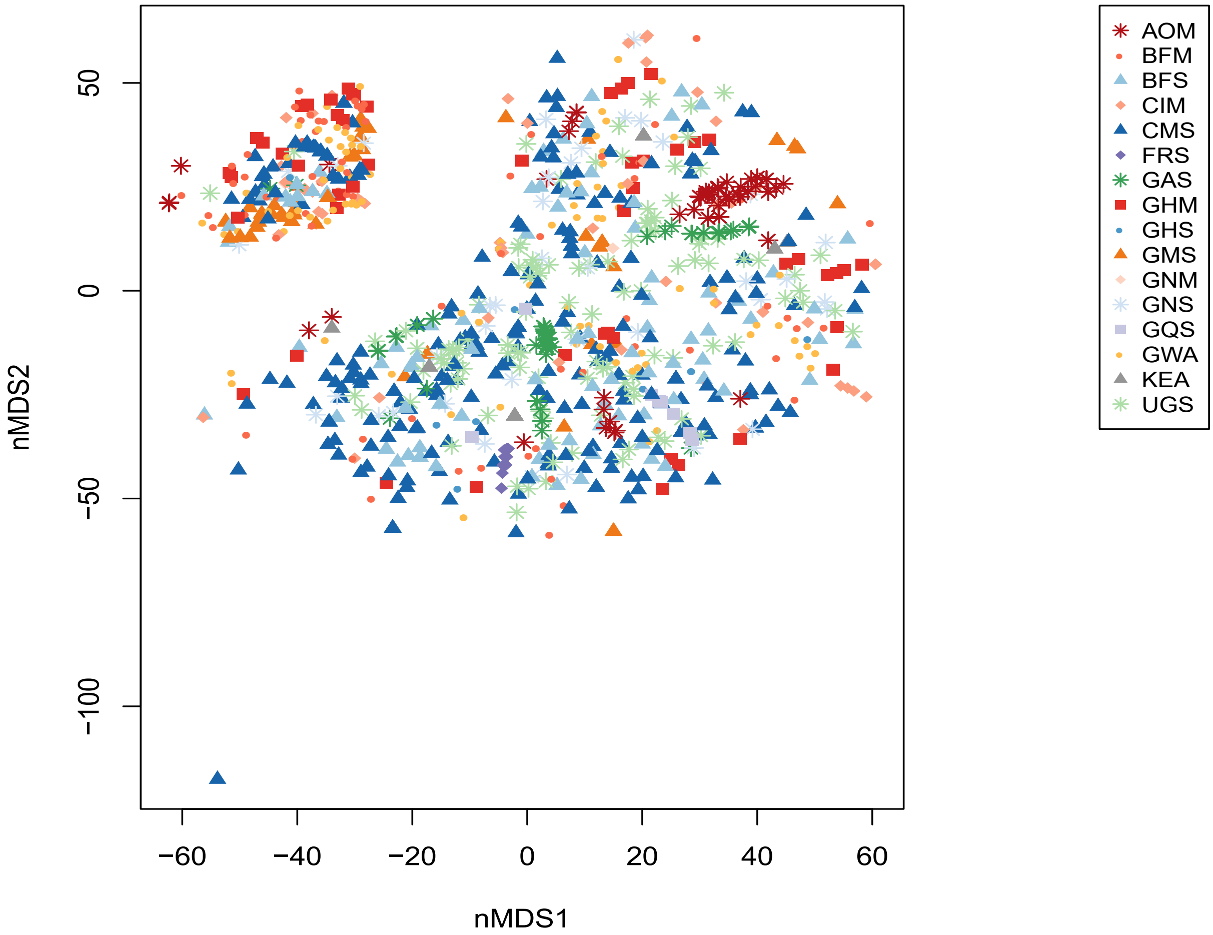


Figure S6. Non-metric Multidimensional Scaling analysis (NMDS) of the mtDNA genetic distance among samples color-coded by populations. The outlier sample at the bottom shows the relict haplotype discovered in one sample (AN0293_C_CMS, see Table S2) and likely reflect a relict haplotype from the *An. arabiensis* mtDNA gene pool before its total replacement by the one from *An. gambiae* and/or *An. coluzzii*. See Table 1 and S2 for the population codes.

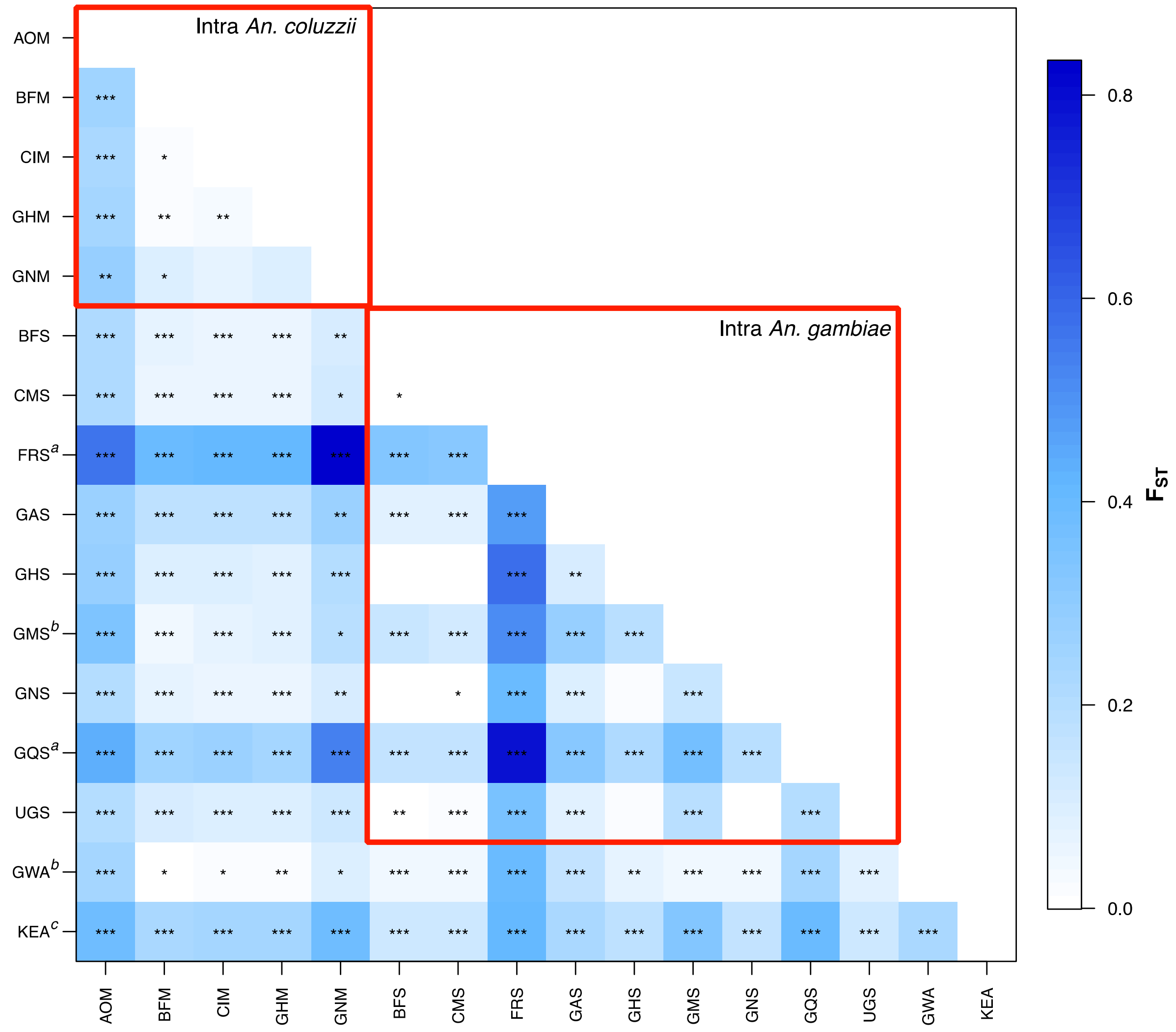


Figure S7. Pairwise level of differentiations expressed as $F_{ST}$ values between populations. (*: p < 0.05; **: p<0.01; ***: p<0.001). Superscripts next to some populations mark (a) the islands populations of Mayotte (FRS) and Bioko (GQS), (b) hybrid taxonomically uncertain populations of The Gambia (GMS) and Guinea-Bissau (GWA), and (c) the taxonomically uncertain population of Kenya (KEA).

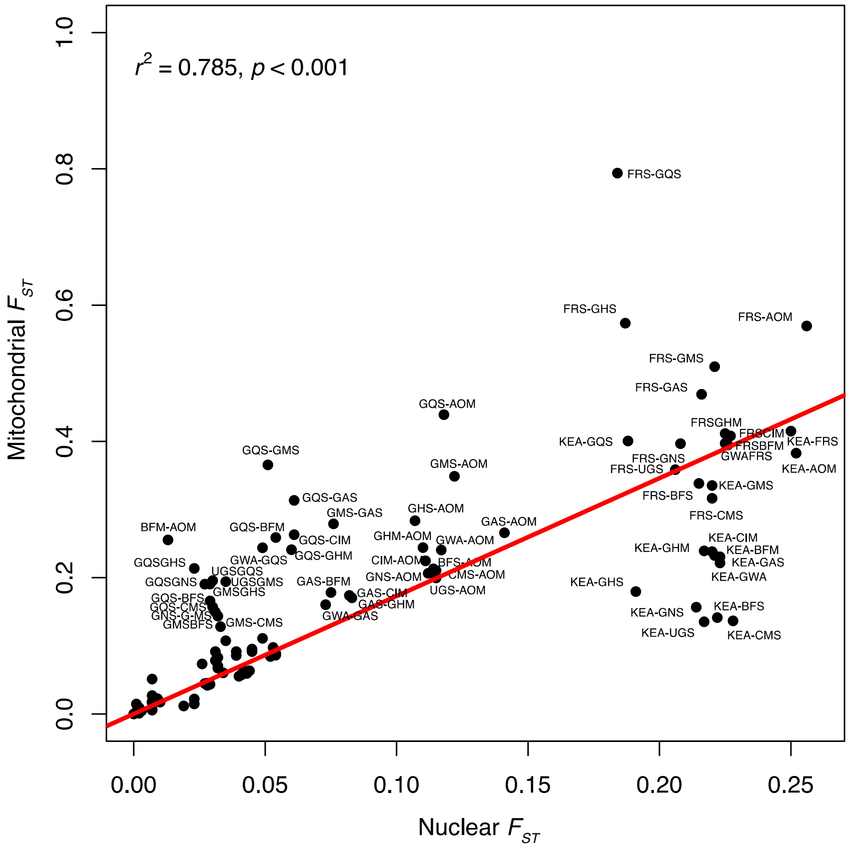


**Figure S8. Linear relationship between the mitochondrial and nuclear $F_{ST}$ values between pairs of populations.** Nuclear $F_{ST}$ values were obtained from The Ag1000G consortium (2020). Pearson's $r^2$ is provided with its associated $p$-value.

*Lenient*

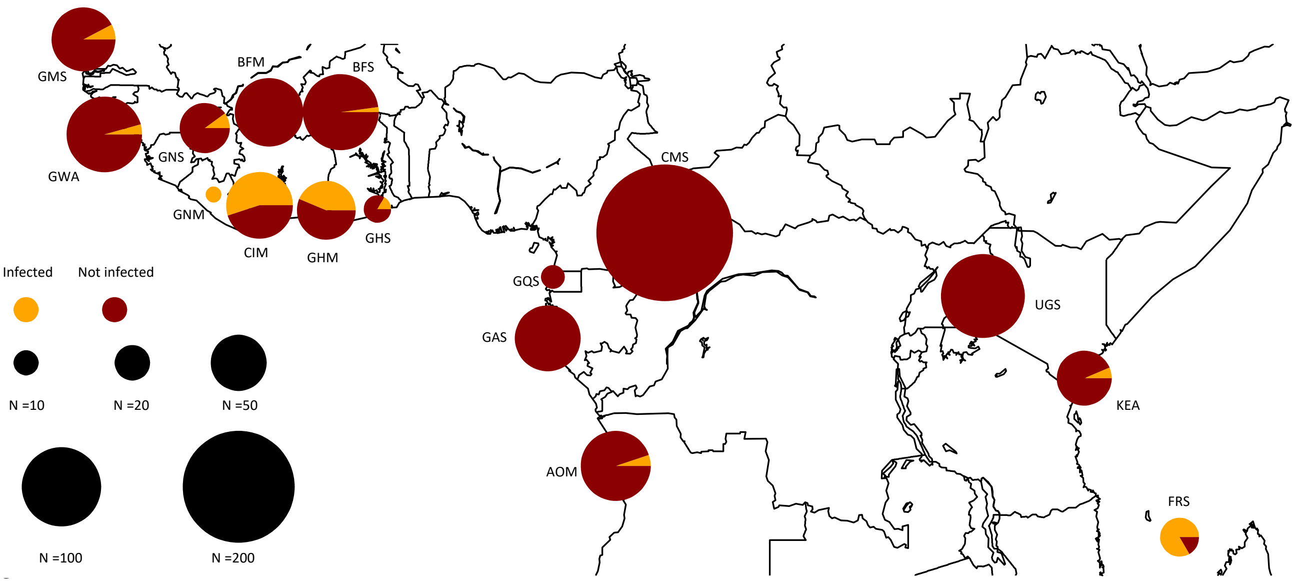


*Strict*

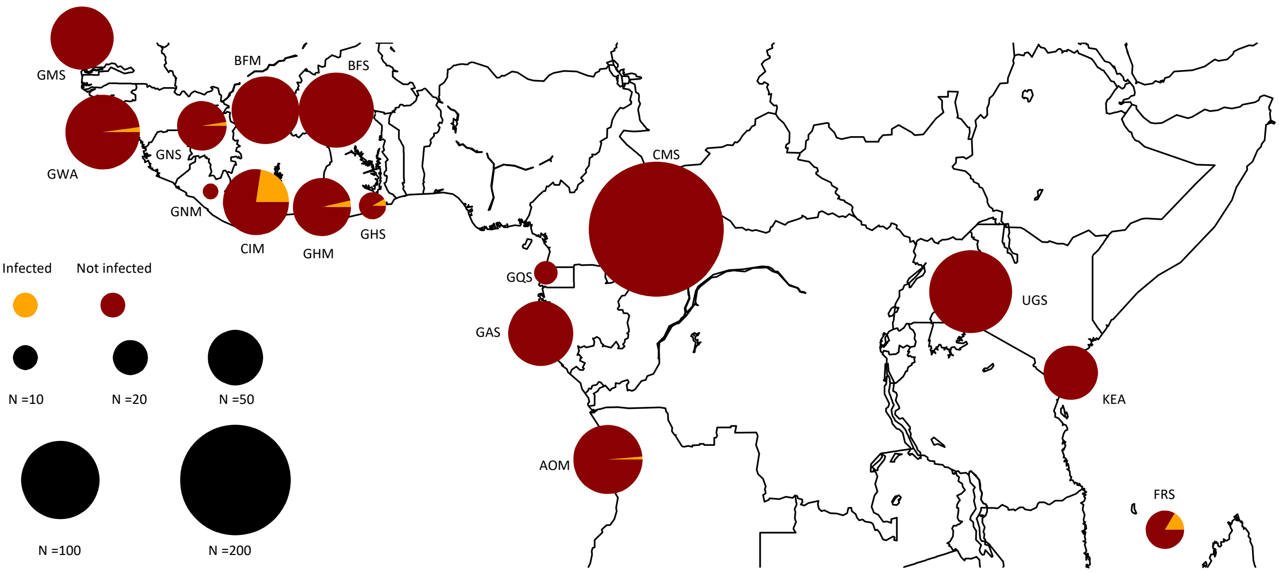


Figure S9. Geographical distribution of rates of *Wolbachia* prevalence in natural population of *An. gambiae* and *An. coluzzii* detected from the unmapped short-data of the The Ag1000G Consortium (2020) using the lenient (top) or strict (bottom) criteria.

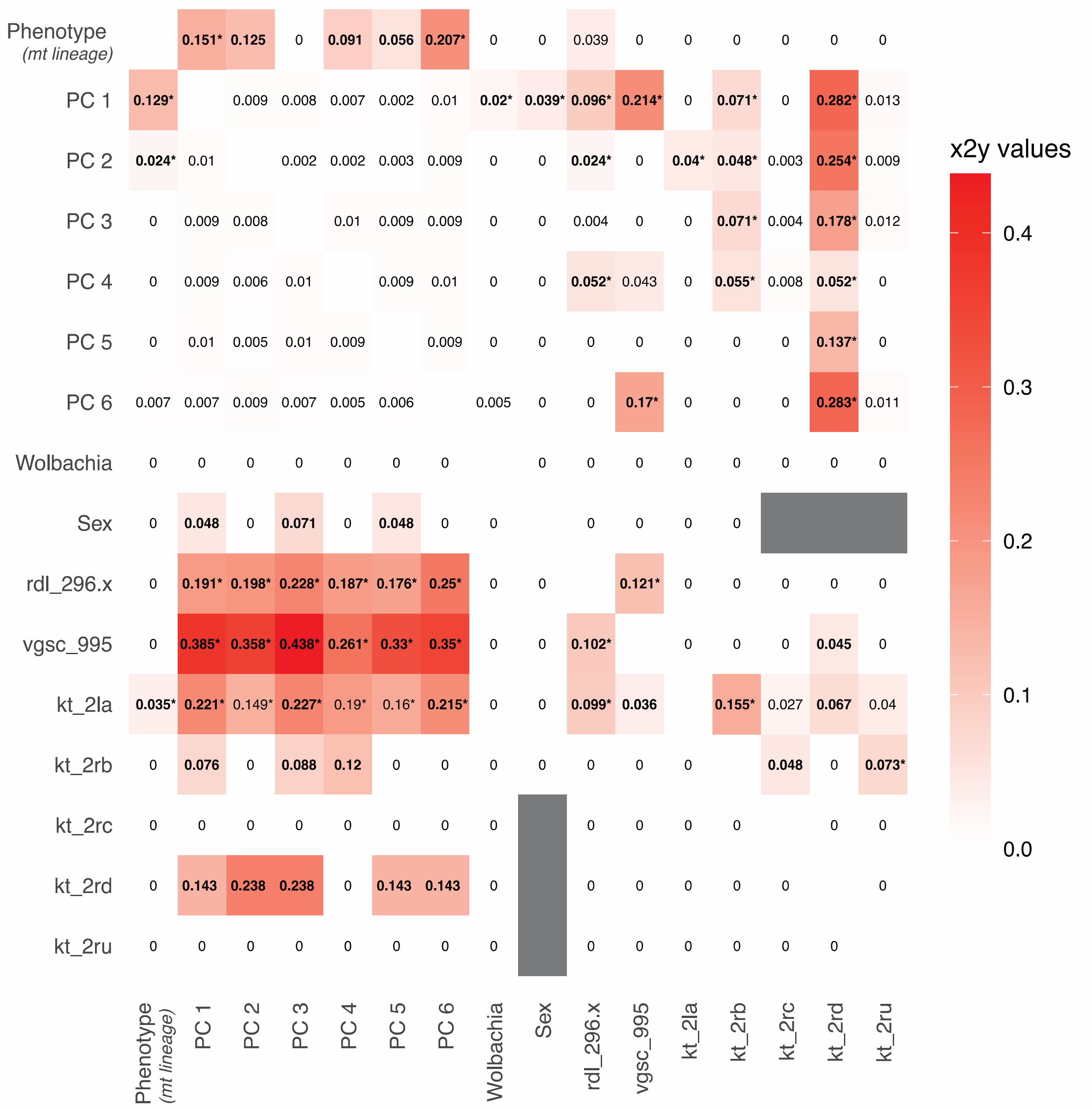


**Figure S10**. Ranked predictive power of cross-features (*x2y*) metric between genomic features. The *x2y* metric is an analogue to the "classic" correlation coefficient for any type of variable (continuous, ordinal, or even qualitative) (Ramakrishnan 2021; Lares 2023). It describes the ability of one variable to predict another one using a CART (Classification And Regression Trees) models. Note that such *x2y* metrics is not symmetric. The extent to which *x* can predict y can be different from the extent to which *y* can predict *x*. Bold values are those explaining more than ≥5% and the bold star values are those for which the 95% CI obtained from 1000 bootstrap resampling excluded zero. The format of the matrix reads as follow: rows (*x*) predicting (~) columns (*y*).

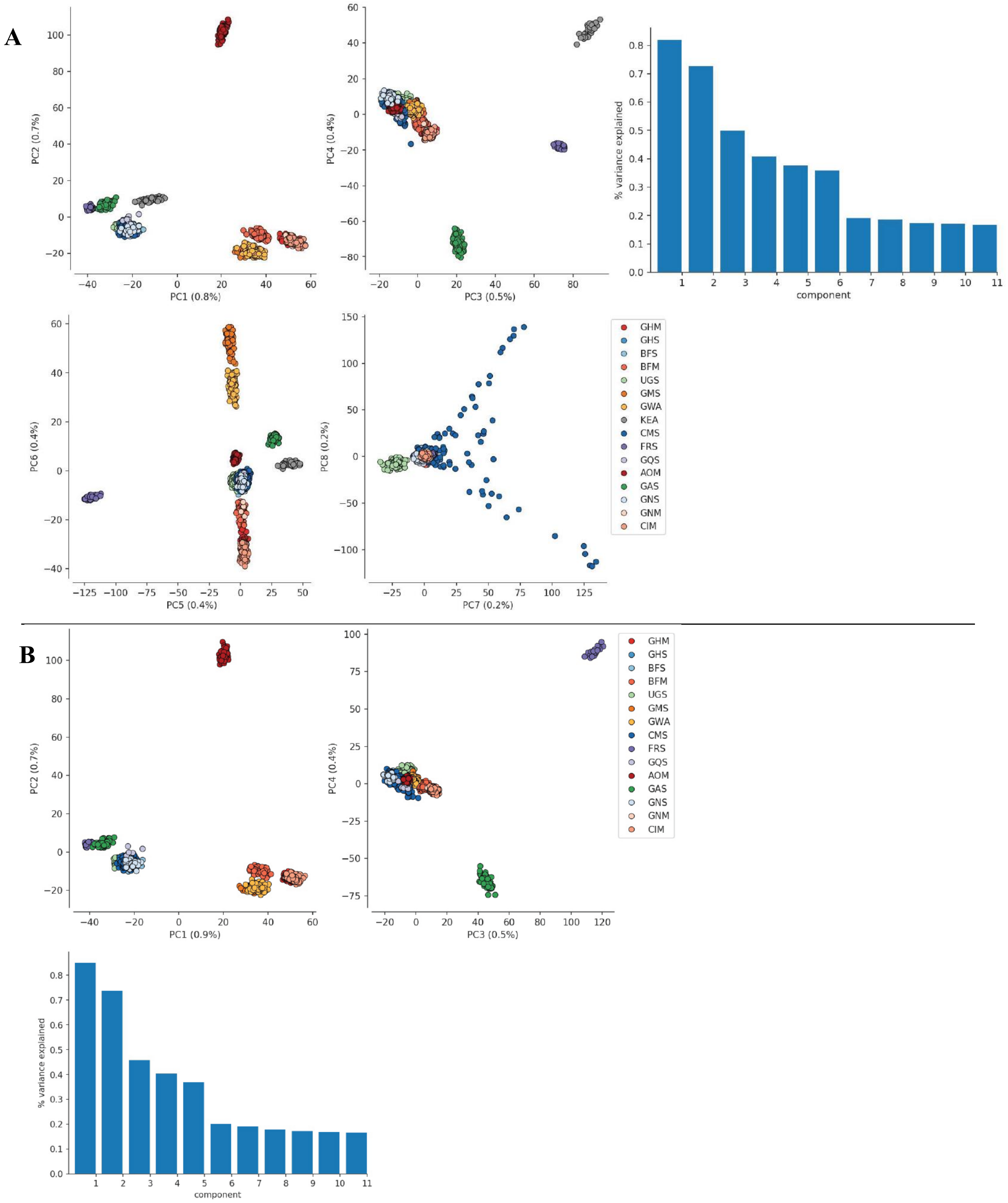


Figure S11. **(A)** Principal component analysis (PCA) describing the population structure of the 1142 samples included in the Ag1000G phase-II consortium project based on the 100,000 unlinked SNPs from the freely recombining regions of chromosome 3L following The Ag1000G Consortium (2017, 2020). PC score values for each sample are shown for PC1 to 8 are shown. Panel **(B)** is the same as panel (A), but without the samples from Kenya. PCs are shown for the 1$^{st}$ tot the 4$^{th}$ PC axes. (See Table 1 and S2 for the population codes).

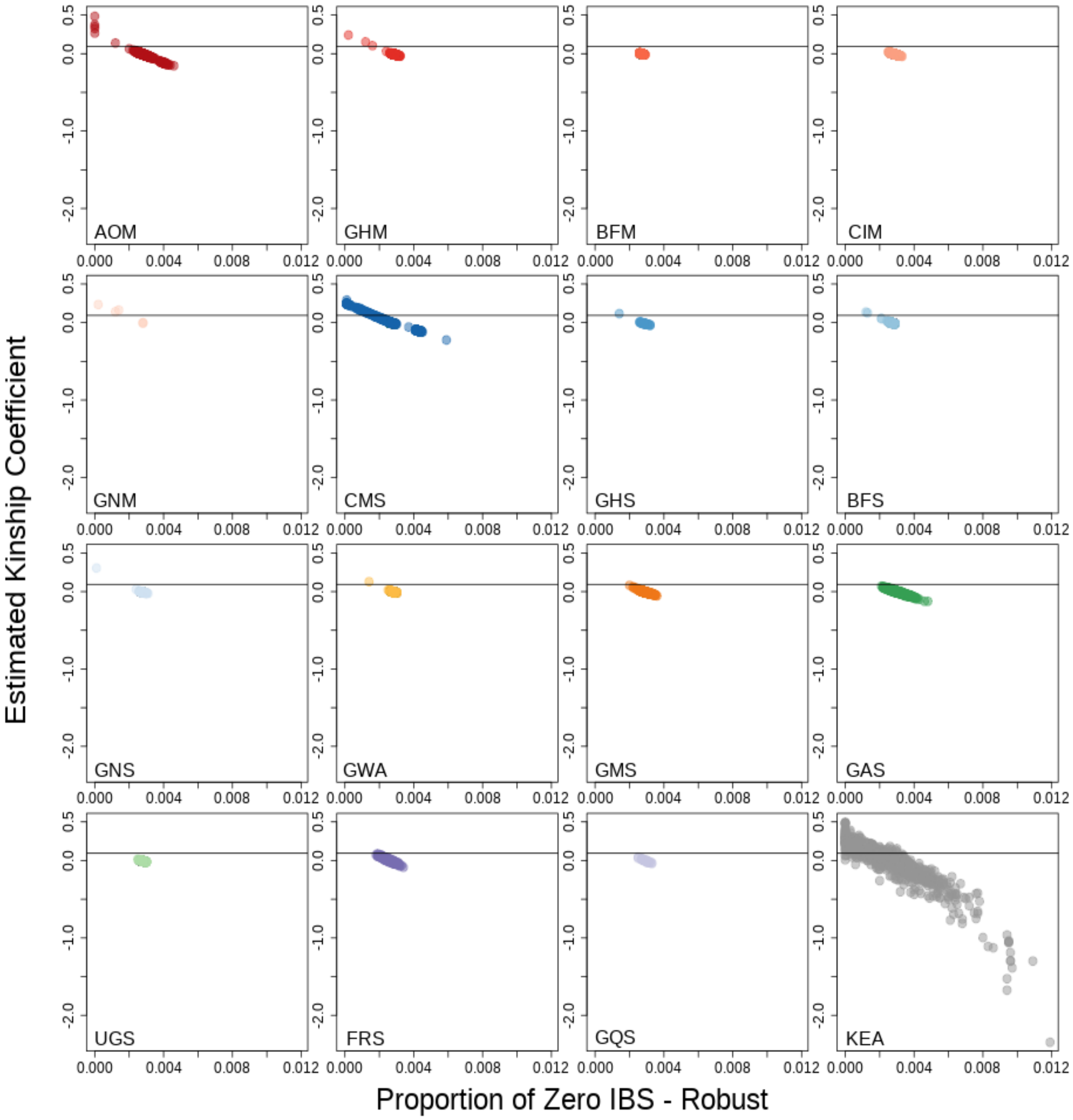


[Figure S12]. Estimated kinship coefficients between pairs of samples, following the KING-robust method which does not assume within population genetic homogeneity, as a function of the probability of zero IBD (or proportion of zero IBS). Zero or negative kinship values represent unrelated samples (see Manichaikul, et al. (2010)). (See Table 1 and S2 for the population codes).

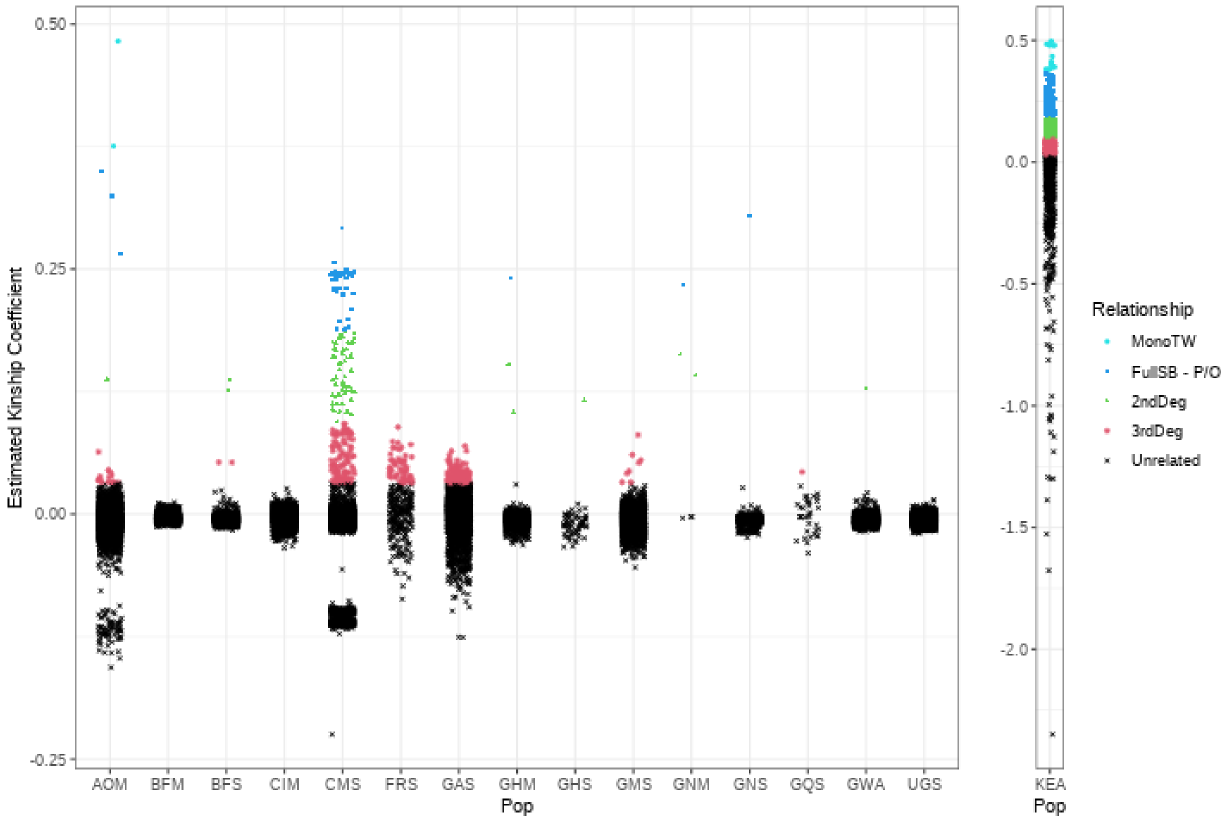


Figure S13. Estimated kinship coefficients between pairs of samples, following the KING-robust method which does not assume within population genetic homogeneity (Manichaikul, et al. 2010). Values were color-coded according to family relationships between pairs of samples according to Manichaikul, et al. (2010) (Table S6). (See Table 1 and S2 for the population codes).

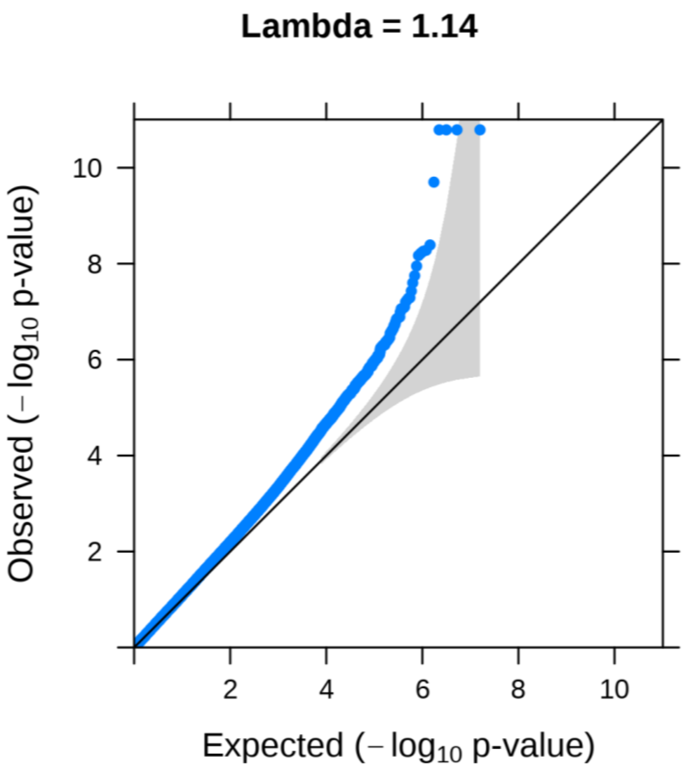


Figure S14. Quantile-to-Quantile plot and a genomic inflation factor (Lambda) close to 1 indicate that the genomic control of the GWAS is globally adequate.

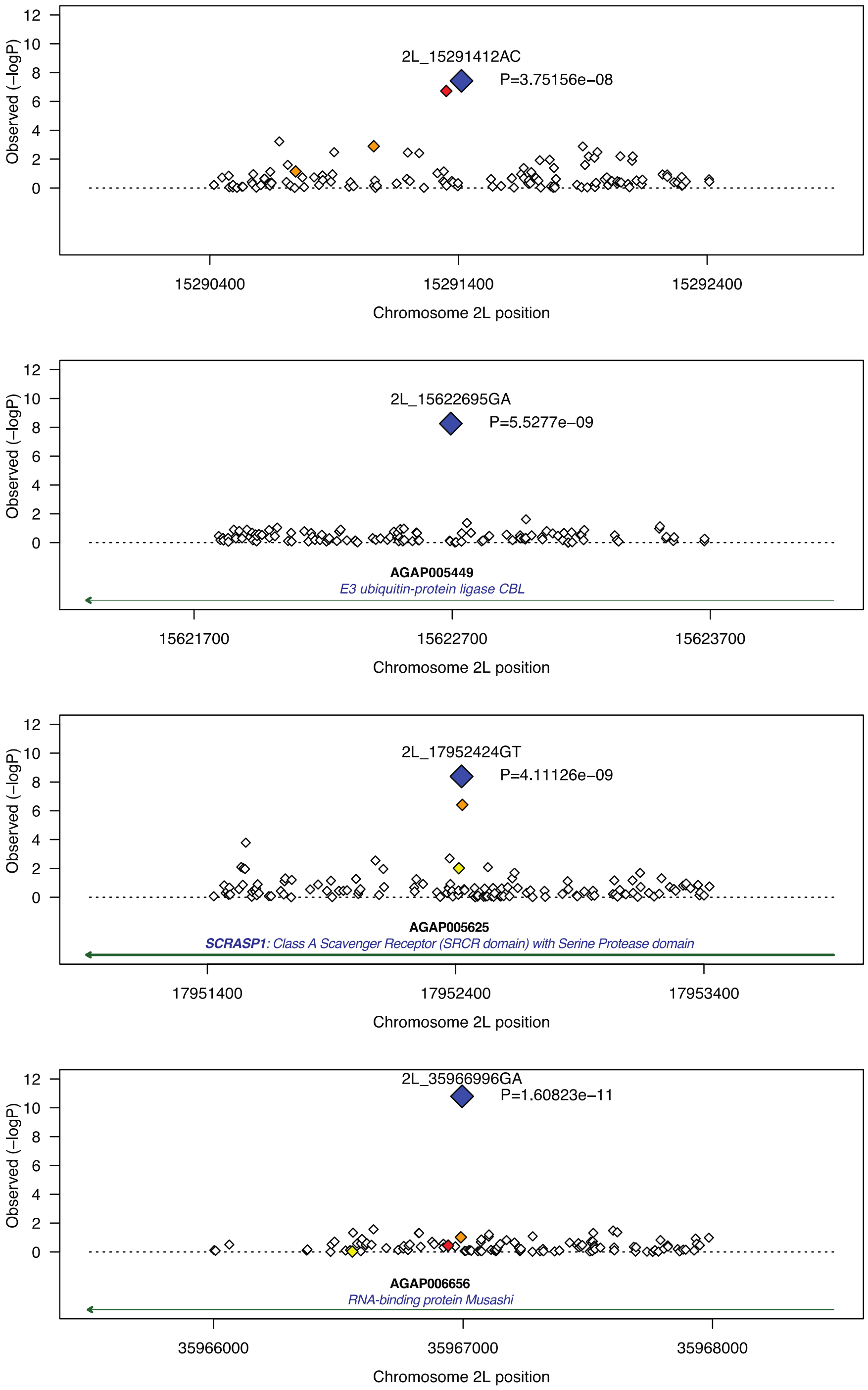


**Figure S15. Signal plot showing the 14 significant SNPs (in blue diamond) identified in the GWAS analysis in perspective with the other SNPs in a 2kb window around it.** The observed *p*-value for each SNP is displayed in $\log_{10}$ scale along the chromosome. The SNPs are color-coded according with LD with between the hit SNP and the other SNPs as follow: SNPs in strong LD are in red ($r^2$ > 0.8); those in moderate LD in orange (0.5 > $r^2$ >0.8); those in weak LD are in yellow (0.2 > $r^2$ >0.5), and the other unlinked SNPs ($r^2$ < 0.2) are in white. The name of the hit SNP is provided as chromosome_ position, the *p-value* estimated in the GWAS analysis, and the gene names are also provided.

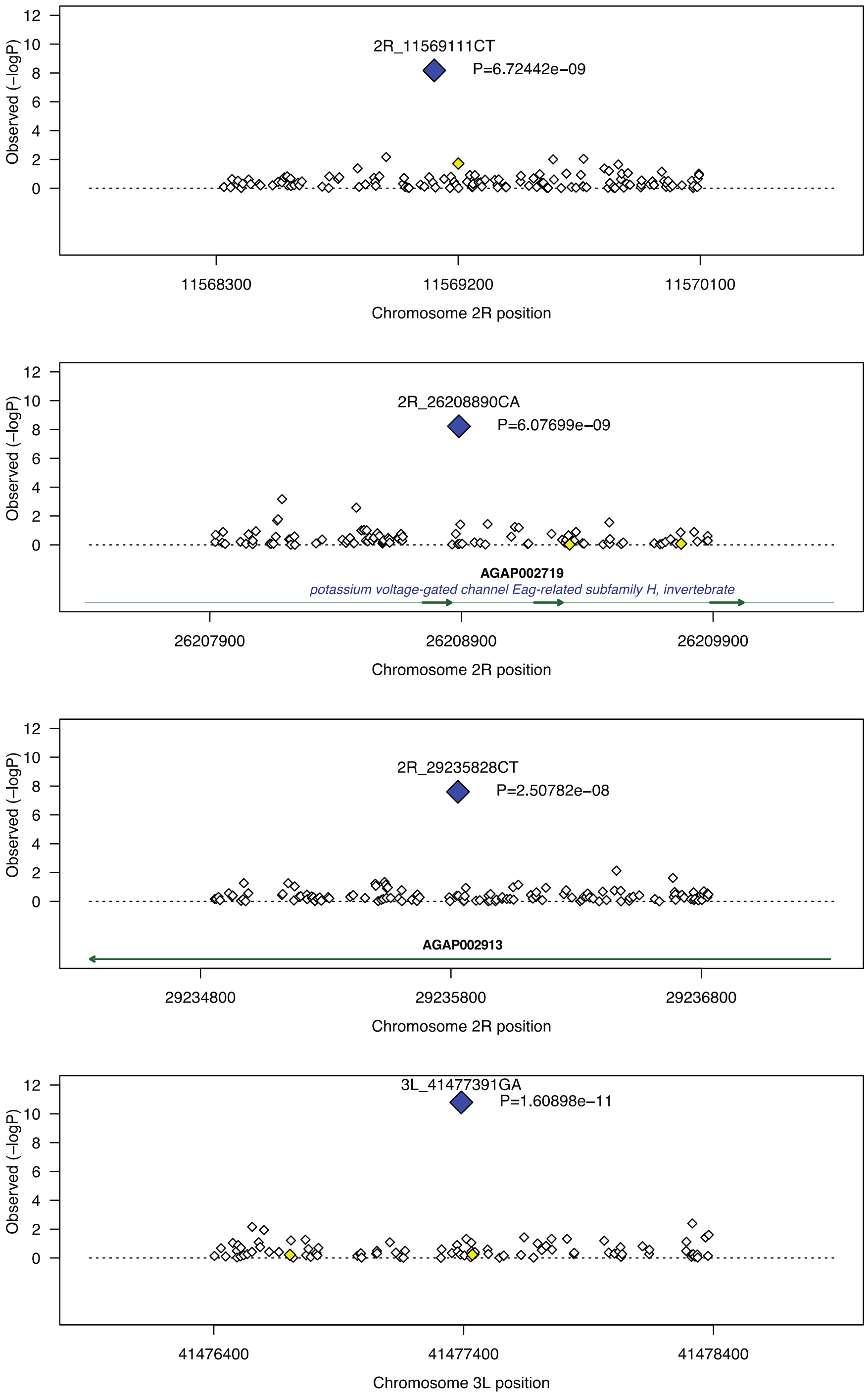


Figure S15. (Continued)

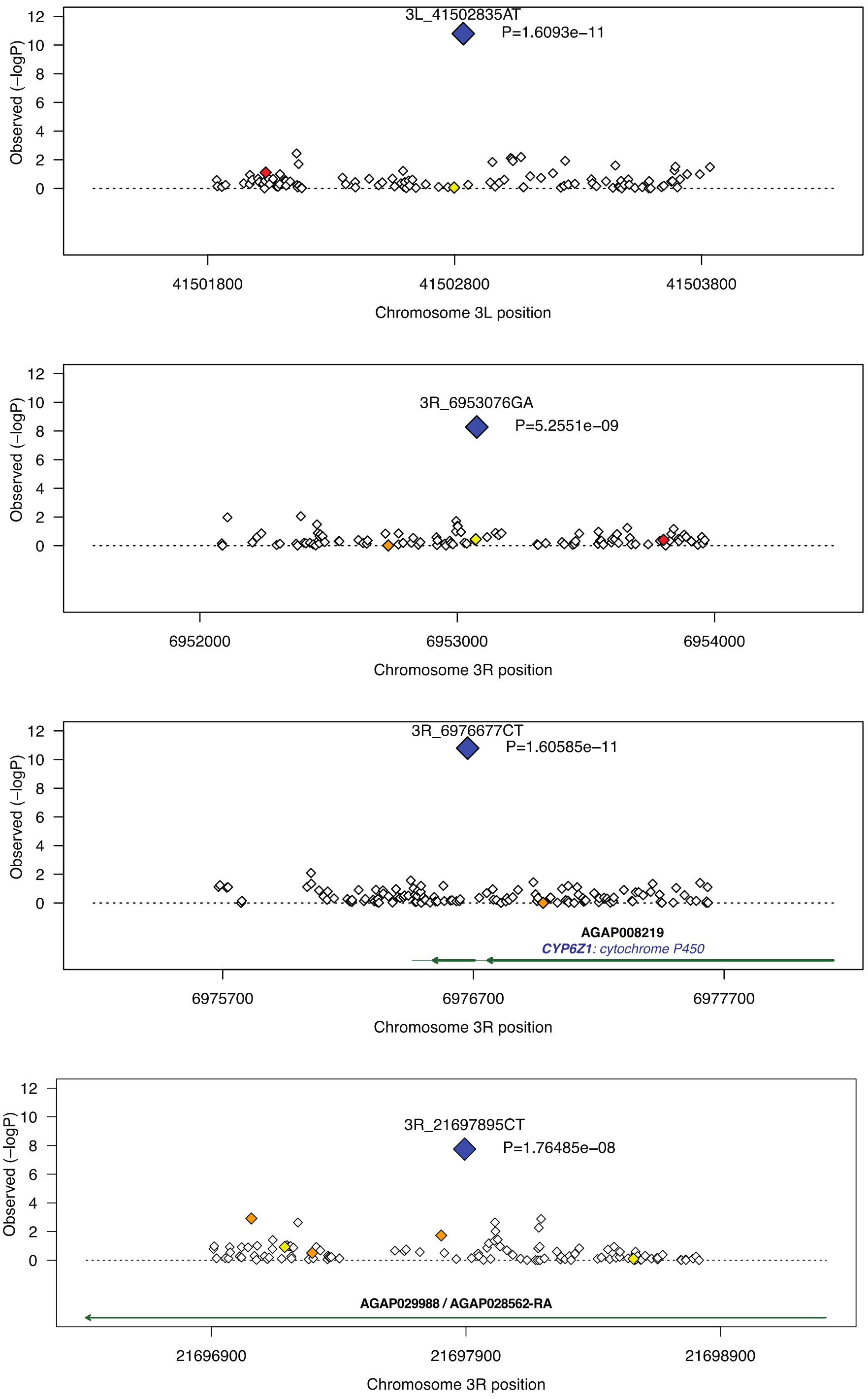


Figure S15. (Continued)

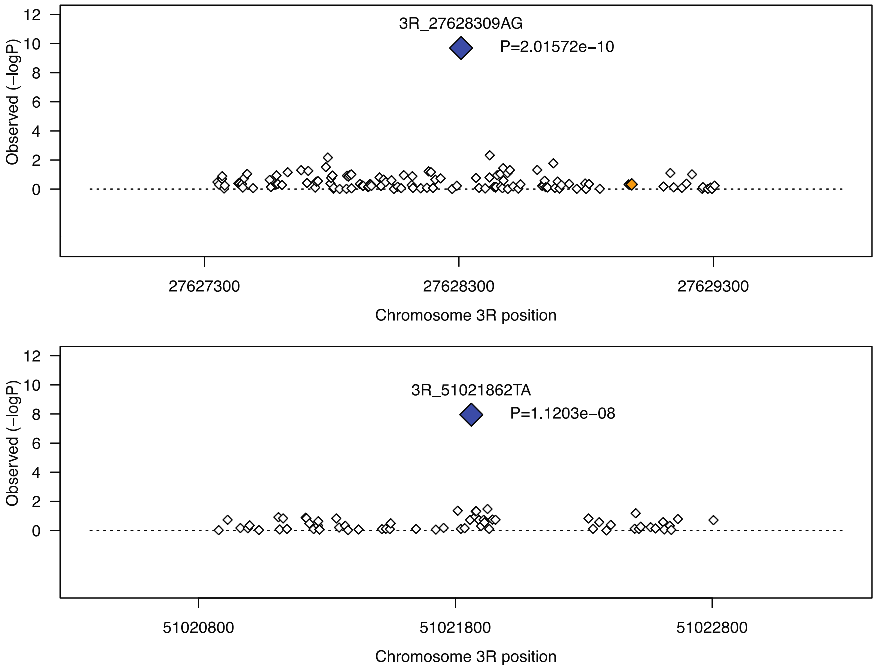


Figure S15. (Continued)

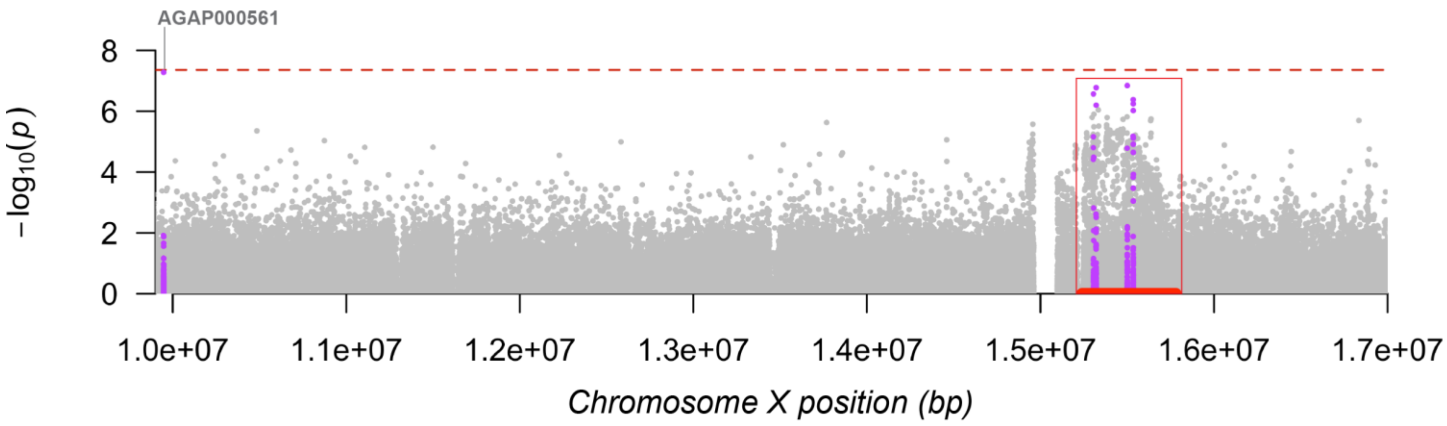


- **AGAP000818**[1] (CYP9K1 - cytochrome P450),
- **AGAP000819** (nuclear receptor subfamily 2 group E member (Tailless)),
- **AGAP000820**[4] (CPR125 - cuticular protein RR-2 family 125),
- **AGAP000821, AGAP000822, AGAP000823** (CD81 antigen),
- **AGAP000824** (bone morphogenetic protein 5), **AGAP000825, AGAP000826** (cap-specific mRNA (nucleoside-2'-O-)-methyltransferase 1),
- **AGAP000829** (calpain-15), **AGAP000830** (CASPS7 - short caspase 7), **AGAP000831** (DnaJ homolog subfamily C member 25),
- **AGAP000832** (Derlin-2/3), **AGAP000833** (MIP - myoinhibitory-like peptide),
- **AGAP000834, AGAP000835, AGAP028655, AGAP013506, AGAP000840** (amiloride-sensitive sodium channel, other),
- **AGAP000841** (Ras-related protein Rab-40), **AGAP013101,**
- **AGAP000842**[1] (NADH dehydrogenase (ubiquinone) 1 alpha subcomplex, assembly factor 1),
- **AGAP000843** (cardiolipin synthase), **AGAP000844** (Progestin and adipoQ receptor family member 4), **AGAP000847** (CDC-like kinase),
- **AGAP000848, AGAP000849** (NADH dehydrogenase (ubiquinone) 1 beta subcomplex 1),
- **AGAP000850, AGAP000851**[1] (cytochrome c oxidase subunit 6a, mitochrondrial),
- **AGAP000852** (Small ubiquitin-related modifier), **AGAP000853** (gamma-glutamyltranspeptidase), **AGAP000854**

Key to insecticide resistance candidate gene types: [1] metabolic; [2] target-site; [3] behavioural; [4] cuticular.

**Figure S16. Zoomed view on the X chromosome (9.9 to 17Mb) of the Manhattan plot (Fig. 4) showing the genetic associations between the two major mtDNA lineages and each of the SNPs on the nuclear genome.** The red dash horizontal line shows the Bonferroni corrected significance threshold of $4.4\ 10^{-8}$. Three SNPs of interest are highlighted in purple (together with SNPs 1kb upstream or downstream) as they display the lowest *p-value.* Note the region between 15.24Mb and 15.78Mb indicated by a red box. This interval encompasses 2 of the highest association signals and is nested within a broader region displaying a distinctive elevation of the *p*-value. The same core area (in red) were also previously identified by The *Ag1000G consortium* (2017) and in the *Ag1000G Selection Atlas* (*in prep; https://malariagen.github.io/agam-selection-atlas/0.1-alpha3/index.html*) as displaying a strong signal of positive selection in many populations of *An. gambiae* and *An. coluzzii*, as well as association with metabolic insecticide resistance (Ingham, et al. 2021; Lucas, et al. 2023). A total of 32 genes overlap with this focal region and are listed below the GWAS plot with their gene-id, the associated gene product. Gene in purple are discussed in the main text.

### Supplementary tables

(*Some tables are in the Supplementary Excel file because they are too big Amaya_Romero_etal_SUPPLEMENT_TABLES.xlsx*)

Table S1 | Dataset of Fontaine *et al.* (2015). **Species** = Species; **Sample_ID** = Sample_ID; **Population** = population code; **Country** = Country; **Region** = Region; **Latitude** = approximated latitude; **Longitude** = approximated Longitude; **rdl_296.x** = Genotype for the resistance to dieldrin codon within the Gaba gene; **vgsc_995** = Genotype for the knock-down resistance codon within the Vgsc gene; **kt_2la** = Karyotype for the 2La inversion inferred from sequence data (0=homozygous standard; 1=heterozygous; 2=homozygous inverted); **kt_2rb** = Karyotype for the 2Rb inversion inferred from sequence data (0=homozygous standard; 1=heterozygous; 2=homozygous inverted); **Assembly_attempt** = Identifier for the number of attempts necessary to get the best possible Assembly (First = One attempt, sampling rate %5, No ambiguities remain; Posterior = Two or more attempts, sampling rate >5%, No ambiguities remain; Best = Two or more attempts, sampling rate =>5%, Some ambiguities remain); **Assembly_rate** = Assembly rate; **Format_of_origion** = WG-SRA format (BAM or FASTQ); **Number_of_reads_used** = Number of sampled reads; **Number_of_read_original_file** = Number of reads in the original file; **Final_number_of_reads_used_in_the_assembly** = Actual reads used by the pipeline to produce the assembly; **Percentage_of_total_reads** = Number_of_read_original_file/ Final_number_of_reads_used_in_the_assembly; **Average_coverage_per_sample** = Average coverage per sample; **Assembly_length_pre_trim** = Assembly length before removing D-LOOP; **GC_content_percent** = GC content of assembly. **Accession ID**.

[Table S1 available in the excel file]

**Table S2. Complete Ag1000G dataset.** **Species** = Species; **Sample_ID** = Sample_ID; **Population** = Population code; **Country** = Country; **Region** = Region; **Latitude** = Approximated latitude; **Longitude** = Approximated longitude; **sex =** sex; **rdl_296.x** = Genotype for the resistance to dieldrin codon within the Gaba gene; **vgsc_995** = Genotype for the knock-down resistance codon within the Vgsc gene; **kt_2la** = Karyotype for the 2La inversion inferred from sequence data (0=homozygous standard; 1=heterozygous; 2=homozygous inverted); **kt_2rb** = Karyotype for the 2Rb inversion inferred from sequence data (0=homozygous standard; 1=heterozygous; 2=homozygous inverted); **kt_2rc =** Karyotype for the 2Rc inversion inferred from sequence data (0=homozygous standard; 1=heterozygous; 2=homozygous inverted; None = Undetermined); **kt_2rd =** Karyotype for the 2Rd inversion inferred from sequence data (0=homozygous standard; 1=heterozygous; 2=homozygous inverted; None = Undetermined); **kt_2rj =** Karyotype for the 2Rj inversion inferred from sequence data (0=homozygous standard; 1=heterozygous; 2=homozygous inverted; None = Undetermined); **kt_2rk =** Karyotype for the 2Rk inversion inferred from sequence data (0=homozygous standard; 1=heterozygous; 2=homozygous inverted; None = Undetermined); **kt_2ru =** Karyotype for the 2Ru inversion inferred from sequence data (0=homozygous standard; 1=heterozygous; 2=homozygous inverted; None = Undetermined); **Assembly_attempt** = Identifier for the number of attempts necessary to get the best possible Assembly (First = One attempt, sampling rate %5, No ambiguities remain; Posterior = Two or more attempts, sampling rate >5%, No ambiguities remain; Best =  Two or more attempts, sampling rate =>5%, Some ambiguities remain); **Assembly_rate** = Assembly rate; **Format_of_origion** = WG-SRA format (BAM or FASTQ); **Number_of_reads_used** = Number of sampled reads; **Number_of_read_original_file** = Number of reads in the original file; **Final_number_of_reads_used_in_the_assembly** = Actual reads used by the pipeline to produce the assembly; **Percentage_of_total_reads** = Number_of_read_original_file/Final_number_of_reads_used_in_the_assembly; **Average_coverage_per_sample** = Average coverage per sample; **Assembly_length_pre_trim** = Assembly length before removing D-LOOP; **GC_content_percent** = GC content of assembly; **PCA1 to PCA6 =** Principal Components; **cov1 to cov6 =** Normalized Principal Components; **Wolbachia_infection_status_strict** = Wolbachia infection status strict (0= not infected, 1 = infected); **Wolbachia_infection_status_lenient** = Wolbachia infection status lenient (0= not infected, 1 = infected); **Cryptic_lineage** = sample identified as part of the “cryptic lineage” based on the hierarchical clustering of the nMDS results (0= no, 1=yes).

[Table S2 available in the excel file]

Table S3. Software version information and their associated reference used in this study.

| Tools name | Verion | Reference | Website |
|---|---|---|---|
| *ade4* | v1.7-22 | (Dray and Dufour 2007) | http://pbil.univ-lyon1.fr/ade4/ |
| *adegenet* | v.2.1.10 | (Jombart 2008; Jombart and Ahmed 2011) | https://github.com/thibautjombart/adegenet/wiki |
| *APE* | v5.6-4 | (Paradis, et al. 2004; Paradis and Schliep 2019) | https://emmanuelparadis.github.io |
| *Arlequin* | v3.5 | (Excoffier and Lischer 2010) | http://cmpg.unibe.ch/software/arlequin35/ |
| *AutoMitoG* | | This study | https://github.com/jorgeamaya/automatic_genome_assembly |
| *Biopython* | 1.65 | (Cock, et al. 2009) | https://biopython.org |
| *ecodist* | 2.0.9 | (Goslee and Urban 2007) | |
| *EggLib* | 3.0.0b21 | (De Mita and Siol 2012; Siol, et al. 2022) | https://egglib.org |
| *Geneious Prime 2023.0.1* | Build 2022-11-28 12:49 | (Kearse, et al. 2012) | https://www.geneious.com/prime/ |
| *KING* | 2.2.4 | (Manichaikul, et al. 2010) | https://www.kingrelatedness.com |
| *Lares / X2Y metrics* | 5.2.1.9000 | (Ramakrishnan 2021; Lares 2023) | https://github.com/rama100/x2y<br>https://rviews.rstudio.com/2021/04/15/an-alternative-to-the-correlation-coefficient-that-works-for-numeric-and-categorical-variables/ |
| *Libdiversity C library* | | | https://bioinfo.mnhn.fr/abi/people/achaz/cgi-bin/neutralitytst.c |
| *MagicBlast* | 1.1.5 | (Boratyn, et al. 2019) | https://ncbi.github.io/magicblast/ |
| *MEGA* | V.7 | (Kumar, et al. 2016) | https://www.megasoftware.net |
| *MIRA* | 4.0.2 | (Chevreux, et al. 2004) | http://sourceforge.net/projects/mira-assembler/ |
| *MITObim* | 1.9.1 | (Hahn, et al. 2013) | https://github.com/chrishah/MITObim |
| *MUSCLE* | 3.8.31 | (Edgar 2004) | |
| *Picard* | 2.13.2 | | https://broadinstitute.github.io/picard/ |
| *PLINK* | 1.9 | (Chang, et al. 2015) | https://www.cog-genomics.org/plink/ |
| *Poppr* | v2.9.3 | (Kamvar, et al. 2014) | https://grunwaldlab.github.io/poppr/ |
| *RAxML* | v.8.2.7 | (Stamatakis 2006) | https://github.com/amkozlov/raxml-ng |
| *Samtools* | v.1.9 \| 1.3.1 | (Danecek, et al. 2021) | https://www.htslib.org |
| *scikit-allel* | 1.2.1 | (Miles and Harding 2016) | https://github.com/cggh/scikit-allel |
| *Scikit-learn* | v.0.22.1 | (Pedregosa, et al. 2011) | https://scikit-learn.org/stable/ |
| *SNPtest* | v2.5.4-beta3 | (Marchini, et al. 2007) | https://www.well.ox.ac.uk/~gav/snptest/#introduction |
| *VCFtools* | v.0.1.16 | (Danecek, et al. 2011) | https://vcftools.github.io/index.html |
| *VectorBase* | r57, 2022-APR-21 | (Giraldo-Calderon, et al. 2015) | https://vectorbase.org/vectorbase/app |

Table S4. **Patterns of isolation by distance among populations.** The linearized level of differentiation ($F_{ST}/(1-F_{ST})$) between populations was correlated with their log geographic (great circle) distances using a Mantel's test (10000 permutations). The test was conducted at different levels: (1) including all populations (with the exception of the hybrid taxonomically uncertain populations GWA, GMS, and KEA; (2 and 3) among population within species; and (4 and 5) among population within species but after excluding isolated populations of Angola (AOM) for An. coluzzii, and for An. gambiae, removing Gabon (GAS) and Mayotte Island (FRS).

| **Comparisons** | **N** | **Distance min (km)** | **Distance max (km)** | ***Pearson's r*** | **Mantel p-value** |
|---|---|---|---|---|---|
| All combined | 13 | 10 | 6 454 | 0.35 | 0.003 |
| *An. coluzzii* | 5 | 392 | 3 158 | 0.96 | 0.017 |
| *An. gambiae* | 8 | 375 | 6 454 | 0.32 | 0.095 |
| *An. coluzzii* (no AOM) | 4 | 392 | 961 | 0.46 | 0.167 |
| An. gambiae (no FRS nor GAS) | 6 | 595 | 4 915 | -0.18 | 0.617 |

Table S5. Wolbachia infection count and percentage per population of the The Ag1000G Consortium (2020). Lenient (LC) and strict (SC) criteria. See Table 1 and S2 for the population code

| Population | Total | Infected (LC) | infected (LC, %) | Infected (SC) | infected (SC, %) |
|---|---|---|---|---|---|
| **AOM** | 78 | 4 | 5.1% | 1 | 1.3% |
| **BFM** | 75 | 0 | 0.0% | 0 | 0.0% |
| **BFS** | 92 | 2 | 2.2% | 0 | 0.0% |
| **CIM** | 71 | 39 | 54.9% | 16 | 22.5% |
| **CMS** | 297 | 0 | 0.0% | 0 | 0.0% |
| **FRS** | 24 | 20 | 83.3% | 4 | 16.7% |
| **GAS** | 69 | 0 | 0.0% | 0 | 0.0% |
| **GHM** | 55 | 24 | 43.6% | 2 | 3.6% |
| **GHS** | 12 | 2 | 16.7% | 1 | 8.3% |
| **GMS** | 65 | 5 | 7.7% | 0 | 0.0% |
| **GNM** | 4 | 4 | 100.0% | 0 | 0.0% |
| **GNS** | 40 | 4 | 10.0% | 1 | 2.5% |
| **GQS** | 9 | 0 | 0.0% | 0 | 0.0% |
| **GWA** | 91 | 4 | 4.4% | 2 | 2.2% |
| **KEA** | 48 | 3 | 6.3% | 0 | 0.0% |
| **UGS** | 112 | 0 | 0.0% | 0 | 0.0% |
| **Total** | 1142 | 111 | 9.7% | 27 | 2.4% |

Table S6. Category of relatedness per kinship coefficient values as described respectively in KING (Manichaikul, et al. 2010) user note.

| Category | KING |
|---|---|
| Duplicate or Monozygotic Twin | KC ≥ 0.375 KC $(\lim 0.5)$ |
| Full Sibling osr Parent / Offspring | 0.1875 ≥ KC < 0.375 |
| $2^{nd}$ Degree Relative | 0.09375 ≥ KC < 0.1875 |
| $3^{rd}$ Degree Relative | 0.03125 ≥ KC < 0.09375 |
| Unrelated | KC < 0.03125 $(\lim -\infty)$ |

**Table S7. Family relationship inference estimated using the KING-robust method from genome-wide SNPs data between pairs of samples in the Ag1000G sampling.**

**N_SNP**: The number of SNPs that do not have missing genotypes in either of the individual;
**Z0**: Pr(IBD=0) as specified by the provided pedigree data;
**Phi**: Kinship coefficient as specified by the provided pedigree data;
**HetHet**: Proportion of SNPs with double heterozygotes (e.g., AG and AG)
**IBS0**: Proportion of SNPs with zero IBS (identical-by-state) (e.g., AA and GG)
**Kinship**: Estimated kinship coefficient from the SNP data
**Error**: Flag indicating differences between the estimated and specified kinship coefficients (1 for error, 0.5 for warning)
**Relationship**: Inferred relationship type from the Kinship coefficient estimated by KING according to threshold set in Manichaikul et al. (2010) (see Table S6)
**Pop:** population code used in this study (See Fig.1 and Table S2)
**Color:** code of each pop. as a 6 hexadecimal digit number of the form #RRGGBB

[Table S7 available in the excel file]

Table S8. Contingency table of the family relationship between pairs of samples estimated by KING (see Table S7) for each type of relationship and each population. Population codes are detailed in Fig. 1 and Table S2.

| | MonoTW | FullSB - P/O | $2^{nd}$ Degree | $3^{rd}$ Degree | Unrelated | Total |
|---|---|---|---|---|---|---|
| **AOM** | 2 | 3 | 1 | 15 | 2 982 | 3 003 |
| **BFM** | 0 | 0 | 0 | 0 | 2 775 | 2 775 |
| **BFS** | 0 | 0 | 2 | 2 | 4 182 | 4 186 |
| **CIM** | 0 | 0 | 0 | 0 | 2 485 | 2 485 |
| **CMS** | 0 | 48 | 70 | 104 | 43 734 | 43 956 |
| **FRS** | 0 | 0 | 0 | 56 | 220 | 276 |
| **GAS** | 0 | 0 | 0 | 132 | 2 214 | 2 346 |
| **GHM** | 0 | 1 | 2 | 0 | 1 482 | 1 485 |
| **GHS** | 0 | 0 | 1 | 0 | 65 | 66 |
| **GMS** | 0 | 0 | 0 | 8 | 2 072 | 2 080 |
| **GNM** | 0 | 1 | 2 | 0 | 3 | 6 |
| **GNS** | 0 | 1 | 0 | 0 | 779 | 780 |
| **GQS** | 0 | 0 | 0 | 1 | 35 | 36 |
| **GWA** | 0 | 0 | 1 | 0 | 4 094 | 4 095 |
| **KEA** | 15 | 211 | 266 | 168 | 468 | 1 128 |
| **UGS** | 0 | 0 | 0 | 0 | 6 216 | 6 216 |
| **Total** | 17 | 265 | 345 | 486 | 73 806 | 74 919 |

Table S9. Samples excluded to solve all the pairwise relationships exceeding 2nd degree relative level.

| Species | Sample_ID | Population | Country | Region | Latitude | Longitude | sex |
|---|---|---|---|---|---|---|---|
| An.coluzzii | AR0001_C | AOM | Angola | Luanda | -8.821 | 13.291 | F |
| An.coluzzii | AR0015_C | AOM | Angola | Luanda | -8.821 | 13.291 | F |
| An.coluzzii | AR0016_C | AOM | Angola | Luanda | -8.821 | 13.291 | F |
| An.coluzzii | AR0024_C | AOM | Angola | Luanda | -8.821 | 13.291 | F |
| An.coluzzii | AR0042_C | AOM | Angola | Luanda | -8.821 | 13.291 | F |
| An.coluzzii | AR0072_C | AOM | Angola | Luanda | -8.821 | 13.291 | F |
| An.coluzzii | AA0098_C | GHM | Ghana | Madina | 5.68 | -0.16 | F |
| An.coluzzii | AA0110_C | GHM | Ghana | Madina | 5.68 | -0.16 | F |
| An.coluzzii | AV0040_C | GNM | Guinea | Koundara | 8.5 | -9.417 | F |
| An.coluzzii | AV0042_C | GNM | Guinea | Koundara | 8.5 | -9.417 | F |
| An.gambiae | AB0158_C | BFS | BurkinaFaso | Bana | 11.233 | -4.472 | M |
| An.gambiae | AB0177_C | BFS | BurkinaFaso | Pala | 11.15 | -4.235 | F |
| An.gambiae | AN0008_C | CMS | Cameroon | Mayos | 4.341 | 13.558 | F |
| An.gambiae | AN0010_C | CMS | Cameroon | Mayos | 4.341 | 13.558 | F |
| An.gambiae | AN0015_C | CMS | Cameroon | Mayos | 4.341 | 13.558 | F |
| An.gambiae | AN0019_C | CMS | Cameroon | Mayos | 4.341 | 13.558 | F |
| An.gambiae | AN0020_C | CMS | Cameroon | Mayos | 4.341 | 13.558 | F |
| An.gambiae | AN0085_C | CMS | Cameroon | Gado-Badzere | 5.747 | 14.442 | F |
| An.gambiae | AN0123_C | CMS | Cameroon | Mayos | 4.341 | 13.558 | M |
| An.gambiae | AN0124_C | CMS | Cameroon | Mayos | 4.341 | 13.558 | M |
| An.gambiae | AN0125_C | CMS | Cameroon | Mayos | 4.341 | 13.558 | M |
| An.gambiae | AN0127_C | CMS | Cameroon | Mayos | 4.341 | 13.558 | M |
| An.gambiae | AN0129_C | CMS | Cameroon | Mayos | 4.341 | 13.558 | M |
| An.gambiae | AN0131_C | CMS | Cameroon | Mayos | 4.341 | 13.558 | M |
| An.gambiae | AN0137_C | CMS | Cameroon | Mayos | 4.341 | 13.558 | F |
| An.gambiae | AN0138_C | CMS | Cameroon | Mayos | 4.341 | 13.558 | F |
| An.gambiae | AN0139_C | CMS | Cameroon | Mayos | 4.341 | 13.558 | F |
| An.gambiae | AN0142_C | CMS | Cameroon | Mayos | 4.341 | 13.558 | F |
| An.gambiae | AN0144_C | CMS | Cameroon | Mayos | 4.341 | 13.558 | F |
| An.gambiae | AN0147_C | CMS | Cameroon | Mayos | 4.341 | 13.558 | F |
| An.gambiae | AN0152_C | CMS | Cameroon | Mayos | 4.341 | 13.558 | F |
| An.gambiae | AN0154_C | CMS | Cameroon | Mayos | 4.341 | 13.558 | F |
| An.gambiae | AN0163_C | CMS | Cameroon | Mayos | 4.341 | 13.558 | F |
| An.gambiae | AN0164_C | CMS | Cameroon | Mayos | 4.341 | 13.558 | F |
| An.gambiae | AN0169_C | CMS | Cameroon | Mayos | 4.341 | 13.558 | F |
| An.gambiae | AN0171_C | CMS | Cameroon | Mayos | 4.341 | 13.558 | F |
| An.gambiae | AN0173_C | CMS | Cameroon | Mayos | 4.341 | 13.558 | F |
| An.gambiae | AN0174_C | CMS | Cameroon | Mayos | 4.341 | 13.558 | F |
| An.gambiae | AN0175_C | CMS | Cameroon | Mayos | 4.341 | 13.558 | F |
| An.gambiae | AN0176_C | CMS | Cameroon | Mayos | 4.341 | 13.558 | F |
| An.gambiae | AN0177_C | CMS | Cameroon | Mayos | 4.341 | 13.558 | F |
| An.gambiae | AN0179_C | CMS | Cameroon | Mayos | 4.341 | 13.558 | F |
| An.gambiae | AN0180_C | CMS | Cameroon | Mayos | 4.341 | 13.558 | F |
| An.gambiae | AN0185_C | CMS | Cameroon | Mayos | 4.341 | 13.558 | F |
| An.gambiae | AN0190_C | CMS | Cameroon | Mayos | 4.341 | 13.558 | F |
| An.gambiae | AN0191_C | CMS | Cameroon | Mayos | 4.341 | 13.558 | F |
| An.gambiae | AN0193_C | CMS | Cameroon | Mayos | 4.341 | 13.558 | F |
| An.gambiae | AN0197_C | CMS | Cameroon | Mayos | 4.341 | 13.558 | F |
| An.gambiae | AN0208_C | CMS | Cameroon | Mayos | 4.341 | 13.558 | F |
| An.gambiae | AN0226_C | CMS | Cameroon | Daiguene | 4.777 | 13.844 | M |
| An.gambiae | AN0251_C | CMS | Cameroon | Daiguene | 4.777 | 13.844 | F |
| An.gambiae | AN0256_C | CMS | Cameroon | Daiguene | 4.777 | 13.844 | F |
| An.gambiae | AN0259_C | CMS | Cameroon | Daiguene | 4.777 | 13.844 | F |
| An.gambiae | AN0261_C | CMS | Cameroon | Daiguene | 4.777 | 13.844 | F |

| | | | | | | | |
|---|---|---|---|---|---|---|---|
| An.gambiae | AN0262_C | CMS | Cameroon | Daiguene | 4.777 | 13.844 | F |
| An.gambiae | AN0270_C | CMS | Cameroon | Daiguene | 4.777 | 13.844 | F |
| An.gambiae | AN0284_C | CMS | Cameroon | Daiguene | 4.777 | 13.844 | F |
| An.gambiae | AN0292_C | CMS | Cameroon | Daiguene | 4.777 | 13.844 | F |
| An.gambiae | AN0317_C | CMS | Cameroon | Daiguene | 4.777 | 13.844 | F |
| An.gambiae | AA0061_C | GHS | Ghana | Madina | 5.68 | -0.16 | F |
| An.gambiae | AV0021_C | GNS | Guinea | Koraboh | 9.25 | -9.917 | F |
| Hybrid | AJ0045_C | GWA | Guinea-Bissau | Antula | 11.891 | -15.582 | F |
| Uncertain | AK0060_C | KEA | Kenya | Kilifi-Junju | -3.862 | 39.745 | F |
| Uncertain | AK0062_C | KEA | Kenya | Kilifi-Junju | -3.862 | 39.745 | F |
| Uncertain | AK0065_C | KEA | Kenya | Kilifi-Mbogolo | -3.635 | 39.858 | F |
| Uncertain | AK0066_C | KEA | Kenya | Kilifi-Mbogolo | -3.635 | 39.858 | F |
| Uncertain | AK0067_C | KEA | Kenya | Kilifi-Mbogolo | -3.635 | 39.858 | F |
| Uncertain | AK0068_C | KEA | Kenya | Kilifi-Mbogolo | -3.635 | 39.858 | F |
| Uncertain | AK0070_C | KEA | Kenya | Kilifi-Mbogolo | -3.635 | 39.858 | F |
| Uncertain | AK0073_C | KEA | Kenya | Kilifi-Mbogolo | -3.635 | 39.858 | F |
| Uncertain | AK0074_C | KEA | Kenya | Kilifi-Mbogolo | -3.635 | 39.858 | F |
| Uncertain | AK0075_C | KEA | Kenya | Kilifi-Mbogolo | -3.635 | 39.858 | F |
| Uncertain | AK0076_C | KEA | Kenya | Kilifi-Mbogolo | -3.635 | 39.858 | F |
| Uncertain | AK0078_C | KEA | Kenya | Kilifi-Mbogolo | -3.635 | 39.858 | F |
| Uncertain | AK0079_C | KEA | Kenya | Kilifi-Mbogolo | -3.635 | 39.858 | F |
| Uncertain | AK0080_C | KEA | Kenya | Kilifi-Mbogolo | -3.635 | 39.858 | F |
| Uncertain | AK0081_C | KEA | Kenya | Kilifi-Mbogolo | -3.635 | 39.858 | F |
| Uncertain | AK0082_C | KEA | Kenya | Kilifi-Mbogolo | -3.635 | 39.858 | F |
| Uncertain | AK0085_C | KEA | Kenya | Kilifi-Mbogolo | -3.635 | 39.858 | F |
| Uncertain | AK0086_C | KEA | Kenya | Kilifi-Mbogolo | -3.635 | 39.858 | F |
| Uncertain | AK0087_C | KEA | Kenya | Kilifi-Mbogolo | -3.635 | 39.858 | F |
| Uncertain | AK0088_C | KEA | Kenya | Kilifi-Mbogolo | -3.635 | 39.858 | F |
| Uncertain | AK0091_C | KEA | Kenya | Kilifi-Mbogolo | -3.635 | 39.858 | F |
| Uncertain | AK0092_C | KEA | Kenya | Kilifi-Mbogolo | -3.635 | 39.858 | F |
| Uncertain | AK0093_C | KEA | Kenya | Kilifi-Mbogolo | -3.635 | 39.858 | F |
| Uncertain | AK0095_C | KEA | Kenya | Kilifi-Mbogolo | -3.635 | 39.858 | F |
| Uncertain | AK0096_C | KEA | Kenya | Kilifi-Junju | -3.862 | 39.745 | F |
| Uncertain | AK0098_C | KEA | Kenya | Kilifi-Junju | -3.862 | 39.745 | F |
| Uncertain | AK0100_C | KEA | Kenya | Kilifi-Junju | -3.862 | 39.745 | F |
| Uncertain | AK0102_C | KEA | Kenya | Kilifi-Junju | -3.862 | 39.745 | F |
| Uncertain | AK0103_C | KEA | Kenya | Kilifi-Junju | -3.862 | 39.745 | F |
| Uncertain | AK0104_C | KEA | Kenya | Kilifi-Junju | -3.862 | 39.745 | F |
| Uncertain | AK0105_C | KEA | Kenya | Kilifi-Junju | -3.862 | 39.745 | F |
| Uncertain | AK0106_C | KEA | Kenya | Kilifi-Junju | -3.862 | 39.745 | F |
| Uncertain | AK0107_C | KEA | Kenya | Kilifi-Junju | -3.862 | 39.745 | F |
| Uncertain | AK0108_C | KEA | Kenya | Kilifi-Junju | -3.862 | 39.745 | F |
| Uncertain | AK0109_C | KEA | Kenya | Kilifi-Junju | -3.862 | 39.745 | F |
| Uncertain | AK0116_C | KEA | Kenya | Kilifi-Mbogolo | -3.635 | 39.858 | F |

Table S10. Number (#) of SNPs per chromosome used in the GWAS analysis, after removing related individuals identified in the Kinship analysis (Table S9)

| Chromosome | #SNPs before MAF | # SNPs removed | #SNPs after MAF | Independent SNPs |
|---|---|---|---|---|
| **2L** | 8 906 423 | 7 233 176 | 1 673 247 | 129 355 |
| **2R** | 12 047 846 | 9 626 669 | 2 421 177 | 217 329 |
| **3L** | 7 897 666 | 6 582 290 | 1 315 376 | 277 384 |
| **3R** | 10 752 701 | 8 877 351 | 1 875 350 | 366 374 |
| **X** | 4 472 265 | 3 898 840 | 573 425 | 148 610 |
| **Total** | 44 076 901 | 36 218 326 | 7 858 575 | 1 139 052 |

**Table S11. SNPs identified as significantly associated in the GWAS.**

SNPs location on the chromosome (CHR), its exact position (Position), the nature of the SNP (SNP), the associated *p-value*, Gene-id of the genes overlapping with the SNPs or within 1 kb upstream or downstream the focal position. Are also provided the Gene name, its annotated function, the Flybase corresponding ortholog(s), the VectorBase link.

[Table S11 available in the excel file]
